# In tissue spatial single-cell metabolomics by coupling mass spectrometry imaging and immunofluorescences

**DOI:** 10.1101/2024.03.22.586317

**Authors:** Landry Blanc, Florent Grelard, Michael Tuck, Véronique Dartois, Antonio Peixoto, Nicolas Desbenoit

## Abstract

In this work, we introduce a multimodal imaging workflow that integrates Matrix-assisted Laser Desorption Ionization Mass Spectrometry Imaging (MALDI-MSI) combined with Immunofluorescence (IF) microscopy to enhance in tissue spatial single-cell metabolomics. The workflow allows to correlate cell populations with associated small molecule distributions by conducting on the same tissue section MSI before IF staining, addressing tissue integrity challenges and joint image analysis.

To process MSI data with IF guidance, we propose an original and advanced computational strategy utilizing Receiver Operating Characteristic (ROC) analysis, allowing to identify ions specific to targeted histological regions based on IF staining. Moreover, in a non-targeted strategy, we introduce a Spatial Coherence Measure (SCM) to distinguish genuine spatial patterns from noise within ion distributions, enhancing spatial metabolomics’ robustness. Then spatial clustering techniques are employed to group ions sharing similar spatial distribution to reveal histological structures, providing complementary insights into metabolite distributions. We validated our workflow mouse spleen section as this organ presents a spatially complex but well-detailed microenvironment.

In conclusion, our multimodal and computational workflow opens new frontiers for diverse biomedical research applications by promoting precise spatial metabolomics in tissue sections.

## Introduction

In the last 30 years, mass spectrometry imaging (MSI) has become one of the most attractive *ex vivo* techniques for the spatial characterization of molecules.^1^ Many fields, such as pharmaceutical,^2^ cancer,^3^ or neurodegenerative disease research,^4^ have profited from the ability of MSI to provide molecular maps of various molecules. The combination of MSI with other image techniques in multimodal imaging workflow has been a growing field of research for a decade,^5,6^ allowing to obtain wider biological and chemical spatial information from tissue samples. Multimodal imaging workflows combining on same tissue section MSI with microscopy techniques have proven valuable, in precisely linking molecular signatures to areas of tissues containing specific types of cells.^7–12^ Recently, single-cell metabolomics has been achieved using MSI with in-vitro cultured cells.^13^ However to our knowledge, spatial single-cell metabolomic/lipidomic in a complex tissue microenvironment was still not reached. Such an approach, offering to fully understand the interplay between cellular identity and metabolite distribution within tissues, faces different challenges. MSI and microscopy acquisitions have to be done on the same tissue section at cellular resolution (below 10μm) to be able to link individual cells to their metabolite content. Metabolite distributions have to be highly preserved during data acquisition, imposing tissue processing techniques that prevent metabolite delocalization. On the other hand, MALDI-MSI scanning conditions at cellular resolution that require high laser power levels need to be optimized to preserve tissue integrity and allow further antibody labeling for identification of different cell populations. Moreover, MSI and Microscopy datasets have to be precisely aligned to match cellular and chemical data. Finally, advanced computational workflows have to be implemented to fully decipher the huge amount of correlative data.

Our approach presents a novel method that combines subcellular resolution MSI (5 µm) compatible with immunofluorescent staining on the same tissue section. This allows for highly precise identification and analysis of individual cells within their tissue context. Additionally, we have developed a robust computational data processing pipeline to integrate these rich datasets, offering a holistic view of the tissue, encompassing both its chemical and histological makeup. To fully derive the benefits of our workflow as well as our original computational strategy, the mouse spleen was chosen as a biological model. This organ plays a role in blood filtration and the immune system. It presents a well-described complex microenvironment, with multiple layers of cells, including different cell types and subtypes.^14^ This histological heterogeneity constitutes an interesting environment to challenge and evaluate our multimodal workflow. Each spleen follicle (white pulp) is organized with a central vein surrounded by T lymphocytes, themselves surrounded by B lymphocytes. Around these follicles, the red pulp contains red blood cells and macrophages, as well as migrating lymphocytes.^15–17^ The junction of white pulp and red pulp constitutes the marginal zone, where various connective tissue cells and immune cell subtypes expressing specific markers can be observed.^15^

To decipher computationally MSI/IF multimodal dataset, we used an open-source Python library, called Esmraldi, dedicated to handling MSI data in multimodal workflow.^18^ We developed a new pipeline on Esmraldi focused on deciphering histological information for multimodal datasets. We demonstrate first the potential of an automated pipeline based on receiver operating characteristic (ROC) analysis to identify ions specific from targeted histological regions,^19^ in opposition to more common univariate statistical tests such as T.test^12,20^ or Mann Whitney U test.^21^ This approach relies on using IF as ground truth to generate binary masks of cell location and performs automatically ROC analysis of all ion’s distributions (called Volcano ROC). The ROC analysis determines the effectiveness of certain ions as classifiers by assessing the overlap of intensity distributions between a specific tissue region and the surrounding tissue, ultimately aiding in the identification of cell biomarkers. Secondly, we introduce an innovative clustering strategy as it relies on ion spatial similarities in opposition to state-of-the-art approaches based on pixel spectral similarities.^8,10,19,22^ To achieve this workflow, we first implement a new measure, the spatial coherence measure (SCM), to select *m/z* presenting relevant spatial distribution. We then used hierarchical clustering to group these ions based on their spatial similarities and revealed different molecular distributions highlighting as much histological area.

The multimodal imaging method presented in this article offers an exciting perspective by reaching ***in situ*** single-cell spatial metabolomics. By coupling MSI with IF on the same tissue section and at high spatial resolution, this approach allows for a paradigm shift in MSI, moving from pixel-centric thinking to a focus on cells, providing a much more precise and detailed view of biological tissues with complex microenvironments. We detailed the meticulous optimization during data acquisition to preserve small molecule distribution as well as the quality of cell immunostaining. For data processing, we use histological information from microscopy and IF to guide MSI analysis and discover molecular signatures. Then, we delve into the molecular distribution provided by MSI to identify spatial patterns that highlight relevant histological areas. This approach has broad applicability across various biological studies. Enabling single-cell spatial omics within tissues holds immense potential for uncovering novel biological insights and furthering our understanding of complex cellular processes.

## Results

### 1. Multimodal acquisition workflow and dataset alignment

To obtain a precise correlation between molecular profiles and cell types for ***in situ*** single-cell spatial metabolomics, MALDI-MSI was performed at a subcellular resolution (down to 5 μm), and both modalities were applied sequentially on a unique tissue section (**Figure 1a**). Consequently, the MSI acquisition was optimized to preserve the quality of the subsequent IF acquisition. Several steps in the sample preparation of MSI can impair IF staining by damaging the IF antibody targets, such as matrix deposition, matrix rinsing, and MSI scan conditions (laser parameters). For this optimization, we have selected antibodies targeting different immune cells commonly found in the spleen: T lymphocyte/CD4, B lymphocyte/B220, and Myeloid cells/CD11b. The results of these preliminary optimization processes are described in the Method section and **Figures S1** and **S2**. Three of the four tested matrices were suitable for this multimodal workflow: 1,5-Diaminonaphthalene (DAN), 2,5-Dihydroxybenzoic acid (DHB), and α-Cyano-4-hydroxycinnamic acid (CHCA). 9-aminoacridine (9AA) is difficult to rinse from tissue and impaired IF acquisition with its strong fluorescence at low wavelengths (**Figure S1 - 9AA panels**). The rinsing solution appears to be a key parameter, as many tested solvents disturb antibody staining. Only acetone allows to keep the staining of all 3 tested antibodies. Finally, laser parameters during MSI acquisition must be tuned to avoid tissue destruction. We observed that limiting its energy to 7 nJ is the best compromise between MSI signal intensity and preservation of IF staining. Ultimately, DAN for negative ionization mode and DHB for positive ionization mode offer the best annotation results based on Metaspace2020 queries. Therefore, subsequent experiments focused on these two matrices (**Figure 1b** and **Figure S2**).

**Figure 1:**
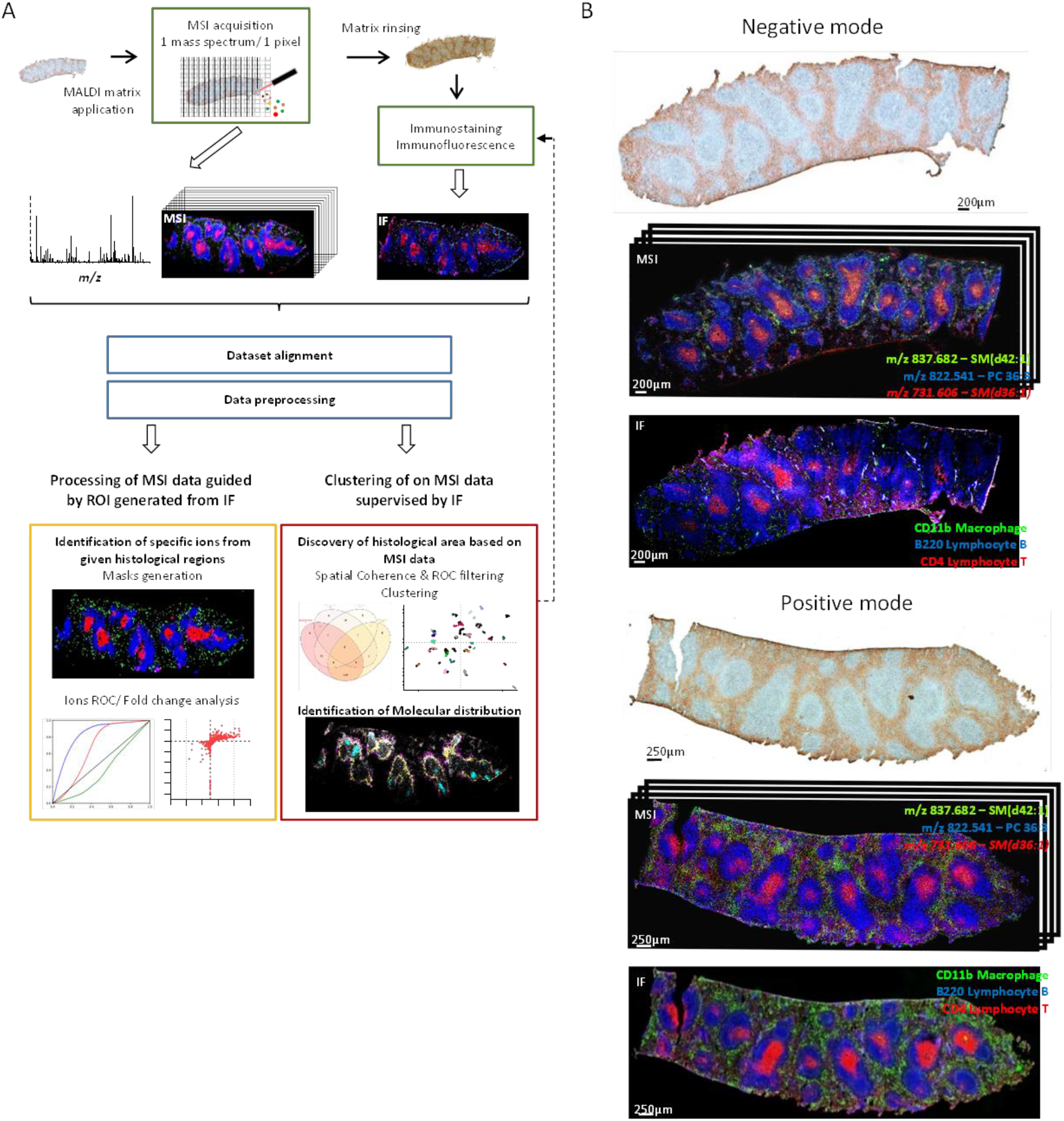
MSI/IF multimodal workflow. Experimental workflow of multimodal acquisition and data processing. B) Spleen sections acquired by MALDI-MSI in negative ion mode with DAN matrix and in positive ion mode with DHB matrix: On top, a bright field image of spleen tissue sections was acquired before matrix application and MSI scan. Middle, MSI data was acquired in negative and positive mode in linear mode and 5um lateral resolution. Bottom, IF staining for CD11b (green), B220 (blue) and CD4 (red) cells receptor. All multimodal datasets are aligned with fiducial marks (see **Figure S3**) Ion annotations made with METASPACE 2020. imzML file for this figure can be found here: https://metaspace2020.eu/dataset/2021-11-09_15h54m35s (for negative mode dataset), https://metaspace2020.eu/dataset/2021-11-08_11h28m30s (for positive mode dataset)

After multimodal acquisition, both datasets are preprocessed. Contrasts of IF images are corrected, and peak selection and alignment, as well as normalization to an internal standard, are done on MS images as described in the Method section. At this stage, both datasets differ in terms of size and spatial orientation. Our optimized MSI acquisition method preserves tissue integrity but does not generate visible ablation marks that could be used to align the two images. Thus, fiducials are etched around the tissue just before MSI acquisition by using a MALDI source laser at full energy (**Figure S3a**). The focalized laser allows the etching of fine marks (around 20μm spots with the shape of a square or cross), which can be used for precise alignment. These marks are visible in the brightfield image, as well as in transmission mode, which is inherently aligned with the IF image. During the etching process, the MALDI matrix and internal standard are fully ablated. Therefore, imprints of those marks are visible as no MSI signal can be observed from them. These marks are then used as fiducial markers to perform an affine registration (**Figure S3b**). The alignment was validated by measuring two metrics: precision and recall, which both evaluate the proportion of pixels in common between the two images. The resulting registration gave precision and recall values higher than 98%, this confirms the high quality of the registration and makes the subsequent analyses possible.

### 2. IF-based histology to guide processing of MSI dataset

To guide the MSI data processing, we selected regions of interest (ROIs) of different cell locations generated within the IF image (**Figure S4a**). First, average MS spectra from relevant areas were extracted (**Figure S4b**). A first insight into the MSI dataset is given by the comparison of average spectra from lymphocyte T in white pulp or in red pulp. Volcano plots were used to automatically find ions that are expressed differently across areas. These display the q-value of a t-test as a function of the fold change. (**Figure S4c**). The results identify metabolites with strong and significant fold change between the two cellular populations. Most of these metabolites are not specific for T cell populations but are significantly more abundant in either white or red pulp, as shown in **Figure S4d**.

To overcome the limitations in terms of specificity of t-test Volcano plots, we use receiver operating characteristic (ROC) curve. In our context, ROC curves reflect the discriminative power of an ion with respect to a binary cell-type mask. Based on microscopy (**Figures 2a** and **b**), we defined different ROIs representing tissue structures or IF-targeted cell populations (binary masks of white pulp, red pulp, B220+ B cells, CD4+ T cells, and CD11b^+^ macrophages, **Figure 2c**). We reduced the resolution of these binary masks (without interpolation) to match the resolution of the MSI dataset. We can perform for each mask a ROC analysis of a given ion, for example, *m/z* 885.549 tentatively assigned as a phosphatidyl-inositol (PI) (38:4) (**Figure** 2d). The area under the curve (ROC AUC) was used to evaluate sensitivity and specificity of an ion with respect to a mask. We also calculated the ratio of mean ion abundance in a mask and its complementary (mask vs off mask).

**Figure 2:**
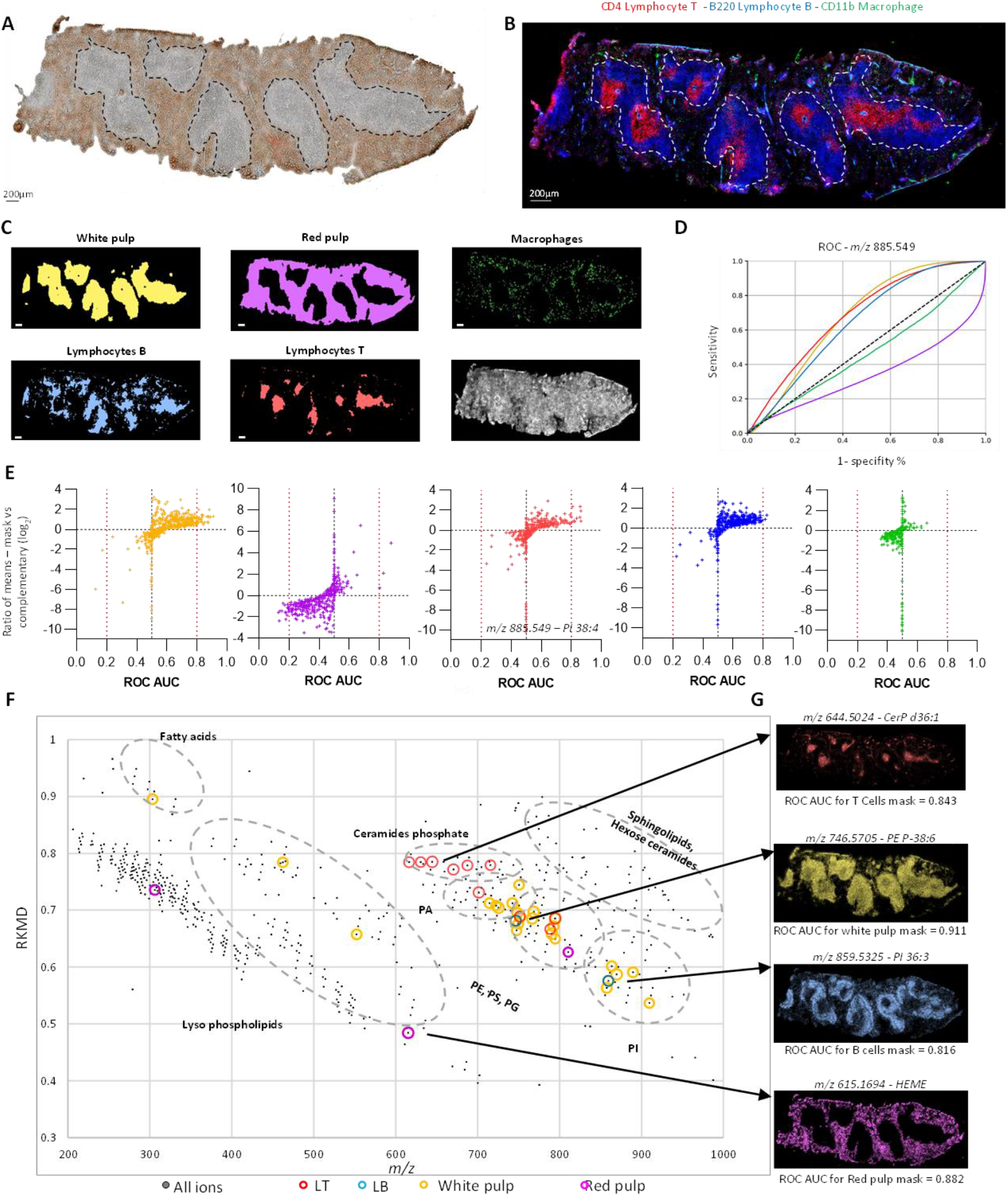
IF to guide MSI data processing: global strategy, multiple ROC analysis – Metabolic fingerprint. A) Bright-field image of spleen tissue section acquired before MSI analysis in negative mode B) IF staining for CD11b (green), B220 (blue) and CD4 (red) cell receptors made after MSI scan. C) IF and brightfield images have been used to generate masks of the location of the different cell populations and tissue structure: White pulp, Red Pulp, lymphocyte T, lymphocyte B and macrophage. MS image of ion *m/z* 885.549 annotated as PI 38:4 is shown as an example. The white scale bar corresponds to 200µm. D) ROC analysis has been used to evaluate the specificity and the sensitivity of ions density map for these masks, here ROC curve for *m/z* 885.549. E) For each ion and each mask, AUC below the ROC curve has been calculated, as well as the ratio of means ion abundance in a mask and its complementary. Similar to a volcano plot, the presented “volcano ROC” plot (ratio of means = f[ROC AUC]), allows a quick visual identification of ions that display large magnitude changes that are also statistically significant for the ROC analysis. F) Kendrick mass defect plot of the list of specific and sensitive ions (AUC of ROC curve >0.8) for each mask. This KMD plot (in base CH_2_) is used organized ions by molecular families. G) MS image of ions with the best ROC AUC for each mask. Ion annotations made with METASPACE 2020. imzML file for this figure can be found here: https://metaspace2020.eu/dataset/2022-04-15_13h38m32s

ROC analyses are automated for each mask and for each peak-picked ion. Then, similar to a volcano plot quickly visualizing ions that display large magnitude changes, we generate a “Volcano ROC” plot: “ratio of means =f(ROC AUC)” (**Figure 2e**). By selecting ions with high ROC AUC (>0.8),^23^ we identify ions with specific and significative accumulation in considered masks (see **Table S1**). Finally, we projected all these ions of interest in a Kendrick mass defect (KMD)^24^ plot (**Figure 2f**) to visualize the whole dataset organized by molecular families. For white pulp, multiple ions from different lipid classes were seen, especially the PE group (**Figure 2g** and **Table S2**). In the red pulp, we highlight two metabolites, glutathione and HEME-^56^Fe. We did not find any ion with high specificity for macrophages, yet we found ions specific for B cells, tentatively assigned phosphatidyl-ethanolamine (PE) P-38:6 (P for a plasmogen lipid side chain) and phosphatidyl-inositol (PI) 36:3. In lymphocyte T, we observed a striking accumulation of ceramide phosphates and PE-ceramide (**Figure 2f** and **g**). Performing a similar volcano ROC analysis in positive mode (**Figure S5**), we identified unique ions for the T cell area. Most of these ions were annotated in Metaspace as different ion adducts of sphingomyelin (SM) d34:1 and d36:1.

Here, we confirm how combining IF and MSI on the same section is a precise tool to spatially study molecular profiles of cell populations. In addition, we show how automatic ROC and “volcano ROC” plots are a useful way to quickly identify molecular biomarkers or to simply highlight metabolite accumulation in a targeted cell population.

### 3. Selection of promising *m/z* to highlight histological areas

MS ion images contain many spatial distributions that are not necessarily explained by regions in IF. For instance, from METASPACE we noticed the distribution of the ion at *m/z* 837.549 strongly located at the border of red pulp and white pulp (**Figure S6**). Such molecular distribution displays a striking organized spatial pattern. We hypothesize that such a distribution highlighted other cell populations or cells with a specific metabolic state and constitute interesting targets to decipher the tissue microenvironment in a molecular histology aspect. To sort the 584 peak-picked ions from the MSI negative mode dataset, we implemented metrics to first automatically eliminate ions that are not specific to the sample area and then to sort the remaining relevant ions based on their spatial coherence. These metrics are non-guided, in the sense that microscopic or histologic masks are not required to sort ion distributions. On the opposite, we implemented also guided measures using histological masks as references of tissue shape.

First, the **off-sample measure** (*OffM*) *e*stimates how points are spread outside of a sample that is centered in the image (**Figure S7a**). We start by estimating an off-sample image by using k-means with two classes and determine for each ion if their signal is mostly off-sample or in-sample. In our metric, values range from 0 to 1, with values close to 0 indicating on-sample ion images. Empirically, we select a threshold of 0.1, below which ion images are considered on-sample (left panel of **Figure 3a**). For the guided approach (right panel of **Figure 3a**), we measure for each *m/z* the ratio of average signal between inside the tissue mask (**Figure S6d**) or its complementary. Positive log2(ratio) or close to 0 indicates a main signal off-sample or nonspecific. A threshold of log2(ratio) lower than −1 was defined to select *m/z* specific from the tissue. We decided to challenge results from the guided and the non-guided in/off-sample (**Figure 3b** Venn diagram). Both methods disagree with only 15 of 584 ions (around 2.6%), showing the efficacy in identifying tissue-specific and non-specific ions. We excluded these 15 ions from further analysis as none appeared tissue-relevant and kept 224 ions for further analysis (**Figure S8a**).

**Figure 3:**
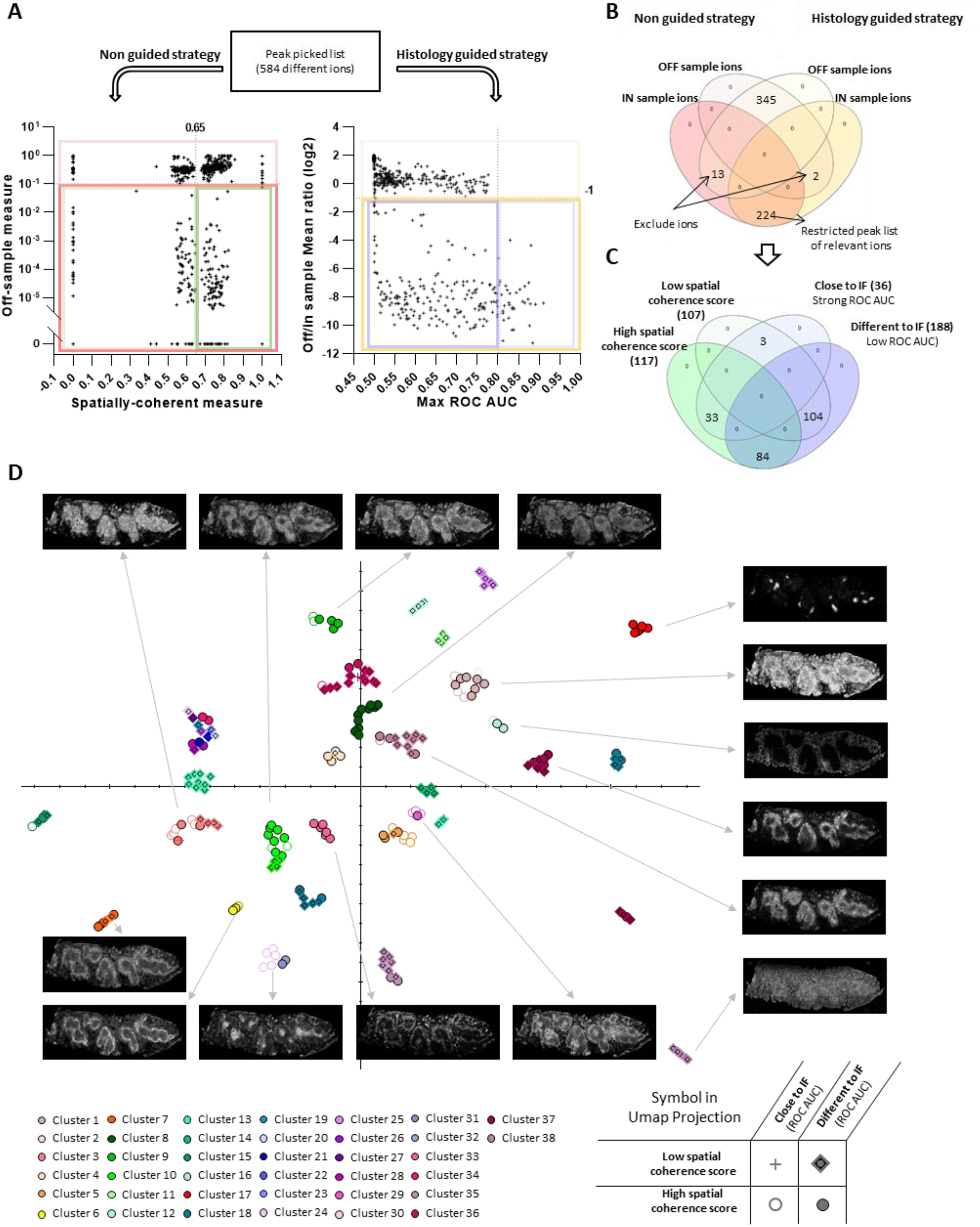
Peaks filtering based on guide and non-guided strategies followed by sorting and clustering based on spatial distribution. A) Partitioning plot obtained for the two strategies. On the left, is the evaluation of ion spatial distributions without any histological input from microscopy or IF. The Off/In measure define automatically where the tissue section is in the image and gives a score for each ion based on whether ion distribution is mostly inside or outside the tissue. The spatial coherence score distinguishes noisy spatial distributions compared to the highly spatially coherent. On the right side, is the evaluation with histologically guided strategy. For each *m/z*, the in/off sample mean ratio is based on the average ion signal inside or outside the whole sample mask (**Figure S6d**), and Max ROC AUC is the best AUC in the ROC curve among the different tested binary masks of the cell population (see **figure 2c**). B) Venn diagram of *m/z* for IN or OFF sample filtering considering the two strategies to select relevant ions. C) Venn diagram to sort relevant m/z based on their spatial coherence and similarities to IF masks. D) UMAP projection of hierarchical clustering of all relevant ions based on their spatial similarities + average images of some spatial clusters.

To detect spatially organized patterns in ion distributions, we propose a new measure, the SCM (**Figure S7b**). The **SCM** selects spatially coherent images by estimating the amount of noise with respect to spatially coherent structures. This measure ranges from 0 to 1, with high values indicating spatially coherent distributions and low values corresponding to noisy ion images. Empirically, we set up a threshold of 0.65, above which ion images are considered spatially coherent. In **Figure S8b**, the efficiency of SCM can be appreciated, with 117 high SCM ions showing sharper spatial structures compared to 107 low SCM ions with noisier structures. In parallel, we implemented a guided strategy relying on IF masks to sort ions based on their ROC AUC observed previously. As shown in **Figure 3c, most of the ions presented a strong AUC in ROC analysis as well as a strong SCM score (33 vs. 3)**, confirming the potential of the SCM as a spatial measure. Interestingly, on the 224 “in sample” ions, a large number (188 in total) do not present a high ROC AUC value whereas 84 of them present a coherent spatial distribution, potentially highlighting interesting histological structure.

### 4. Spatial clustering and spectral clustering: complementary strategies to overview of histological structures highlight by molecular fingerprinting

To highlight histological areas, we implemented a spatial clustering strategy to group ions presenting similar spatial distributions. We used hierarchical clustering along with a projection of the clusters into a reduced space using uniform manifold approximation and projection (UMAP).^25^ We used the cosine metric to build the distance matrix. The number of clusters was chosen with the inconsistency coefficient^26^ of the hierarchical clustering tree (**Figure S7c**) set at a threshold of 1.7. This resulted in 38 clusters (listed in **Figure S8 d, e**, and **f**), which were further visualized by UMAP projections (**Figure 3d**) and modified to separate clusters in the reduced space using the previous distance matrix. The UMAP projection in **Figure 3d** estimates the similarities between ion distribution from each other and identify probable over-clustering (clusters #19 to #24, #26 to #27, and #34) or under-clustering (clusters #35 and #37). A detailed montage of each cluster’s different ion distributions shows how efficient this clustering approach (**Figure S8f**) is to group based on spatial similarities. In addition, we sorted the 224 relevant ions into four distinct groups based on their ROC AUC and SCM scores to evaluate how they spread in the spatial clustering. It appears that ions presenting strong ROC AUC for the same mask can be split into clusters, like good biomarker ions of T.cells split between clusters # 29 and #30 or ions of white pulp between clusters #1, #3, and #5. More strikingly, clusters displaying the sharpest spatial structure, like #6, #17, #29, #30, or #33, exclusively contain ions with high SCM values, reinforcing the interest in this metric.

Finally, we compared this spatial clustering strategy with the more common spectral clustering approach (**Figure S9**).^22^ For spatial clustering, we generated average images of each of these distribution groups to simplify the overview of clusters. For spectral clustering, we used the k-means algorithm^27^ with the same number of clusters found by spatial clustering. Both clustering techniques were compared to identify the commonalities and specificities of each technique. On one hand, some tissue structures are identified by both methods (**Figure 4a**) like spatial cluster #30 and spectral cluster #37, highlighting the T.cells area. On the other hand, some are identified specifically by one or the other methods (**Figure 4b**), like some structures located at the border between red pulp and white pulp highlighted by spatial clusters #6, #20, and #33 or spectral clusters #6.

**Figure 4:**
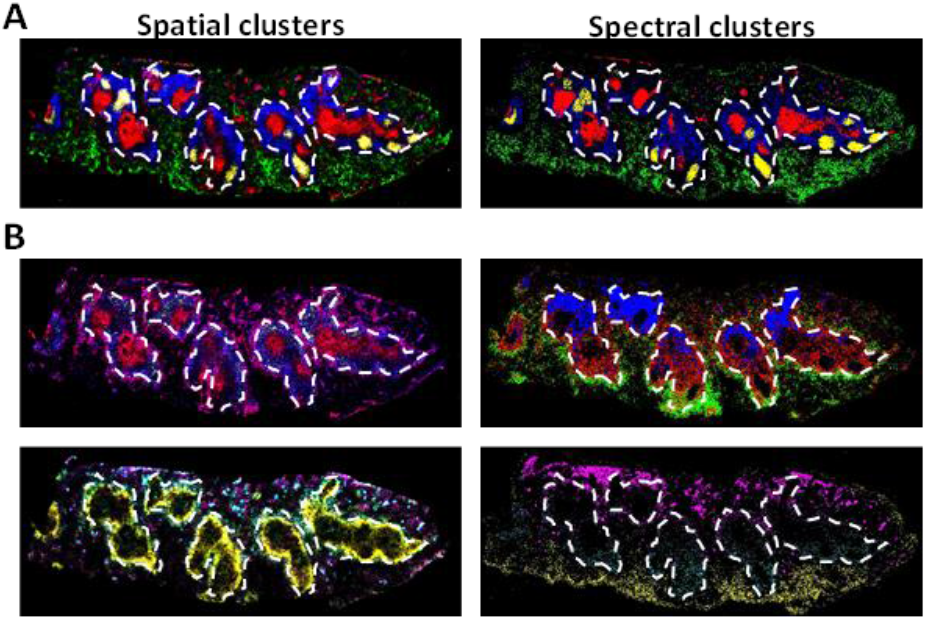
Spatial clustering vs spectral clustering: Complementary strategies to identify different relevant histological areas. Similar histological structures observed from both clustering strategies. On the left, average images of spatial clusters 30 (red), 16 (green), 37 (blue) and 17 (yellow). On the right, spectral clusters 37 (red), 12 (green), 24 (blue) and 36 (yellow). Histological structures highlight only by one of the two the clustering strategies. On the left average images of spatial cluster 29 (red), 20 (green), 7 (blue), 33 (cyan), 34 (magenta) and 6 (yellow). On the right, spectral clusters 1 (red), 6 (green), 13 (blue), 14 (cyan), 34 (magenta), 30 (yellow). White dash line highlights white pulp/red pulp border.

These results indicate that spatial and spectral clustering strategies are complementary approaches, and both should be applied in parallel to highlight histological structures efficiently.

### 5. Identification of histological structure highlighted by MSI

As we identify various ions with consistent and interesting distributions that were previously unseen in IF, we wanted to annotate the associated cell populations, or in other words, select appropriate antibodies to show these cells in IF. We wished to identify cells associated with spatial clusters #6, #20, and #29, localized at the edge of the red pulp and the white pulp.

Candidates for cell populations can be selected by investigating the literature. We focused on the cell population observed at the edge of the white pulp, close to the marginal zone.^15^ We identified three cell populations that could match such distributions: marginal zone macrophages (MZM, positive to CD209),^15,16^ marginal metallophilic macrophages (MMM, positive to CD169)^15–17^ and marginal zone B cells (MZB, positive to CD1d).^16,17^ We selected antibodies targeting these cell populations, cut new spleen sections, and stained them directly without a prior MSI scan, to validate staining in comparison to literature (**Figure S10**). Staining for MZM (CD209) and MMM (CD169) appeared to be closer to the distribution of targeted clusters.

Therefore, we performed a multimodal acquisition of a new spleen section, still with MSI scan first followed by IF staining with new antibodies targeting MMM and MZM (**Figure 5a**). We generated the binary image of the IF signal of CD169 and CD209 for the mask of the MMM (**Figure 5b**) and MZM (**Figure S11a and b**) respectively. We performed a Volcano ROC analysis by comparing binary masks of CD169+ cells (MMM, **Figure 5b**) and CD209+ cells (MZM, **Figure S11b**) with each peak-picked ion (**Figure 5c** and **Figure S11d**). For the MZM mask, none of the peak-picked ions presents relevant ROC AUC (**Figure S11d, e** and **f**). For MMM, however, 3 ions of the spatial cluster#6; m/z 837.549 / 863.569 / 889.581 present ROC AUC >0.85. Ion m/z 837.549 (**Figure 5d**) presents an interesting overlay of MMM in the spleen as shown in **Figure 5e**. These 3 ions have a similar hot spot distribution (**Figure 5d, e**, and **f**) and interestingly they are annotated in METASPACE as part of the same family: respectively PI 34:0 – PI 36:1 – PI 38:2. Therefore, we validated the annotation of these ions by MSMS imaging (**Figure S12**). We confirmed that they are respectively PI 18:0/16:0, PI 18:0/18:1, and PI 18:0/20:2, hinting at a family of molecules, that share similar features in a subclass of macrophage (MMM). We demonstrate that our spatial clustering strategies of molecular distribution can point at specific cell sub-populations, like spatial cluster #6 highlighting the MMM area.

**Figure 5:**
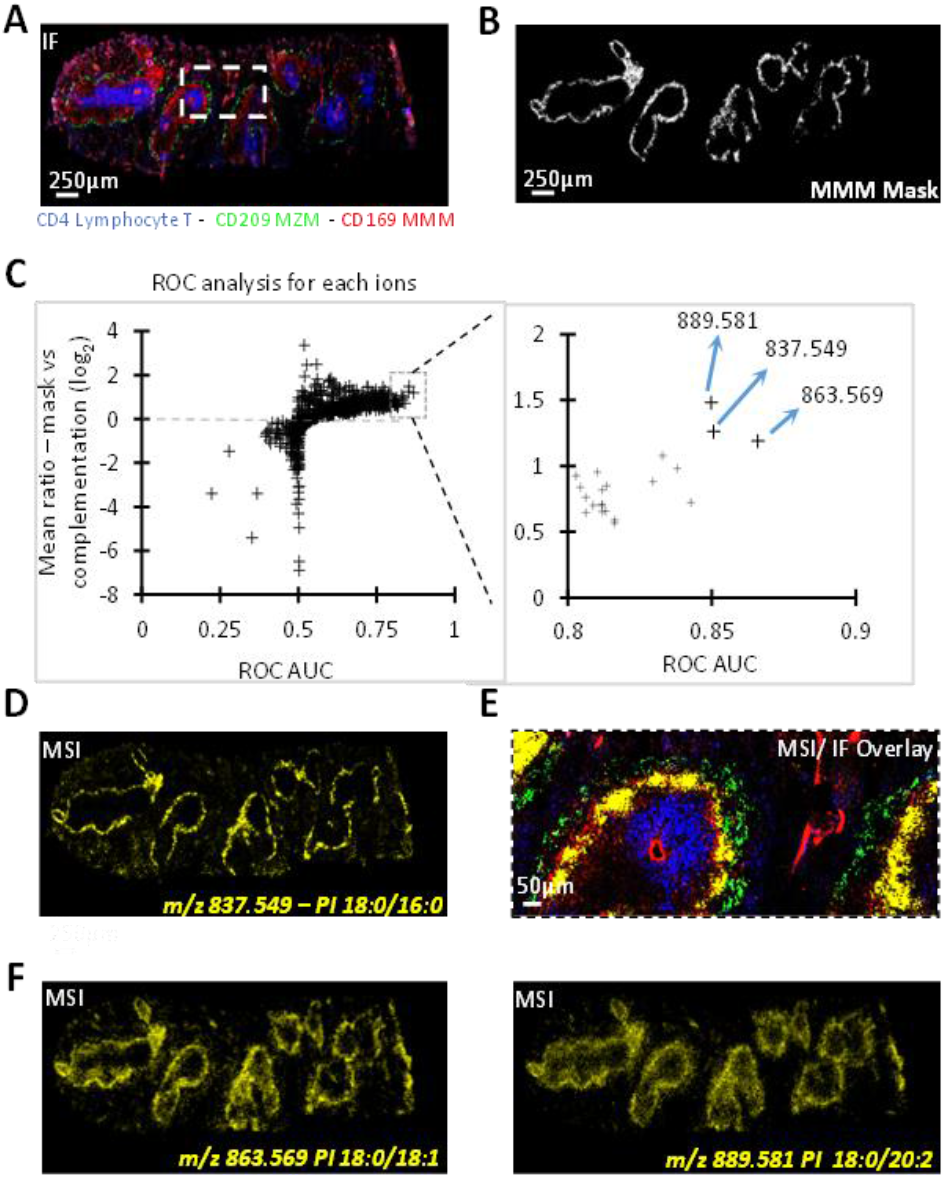
Identification of cell population highlighted by MSI. IF image of the same spleen tissue section stained for CD4, CD209, and CD169 cell receptors. Display of the mask of MMM cells used for ROC analysis. Multiplexed ROC analysis of all detected ions for MMM cells mask. The right panel is a zoom on the best MMM biomarkers, highlighting the 3 best ions (m/z 837.549mm, m/z 863.569, and m/z 889.581, all belonging to spatial cluster#6). D) MSI image of *m/z* 837.549 (PI 18:0/16:0) on a new spleen section. E) Overlay of MSI and IF image. Distribution of m/z 837.549 matches with CD169^+^ MMM cells. F) MSI distribution of ions in the spleen *m/z* 863.569 and *m/z* 889.581. Ion annotation made with METASPACE 2020 and confirmation by MSMS imaging (**Figure S12**). imzML file for this figure can be found here: https://metaspace2020.eu/dataset/2022-04-17_19h49m12s.

## Discussion

Spatial metabolomics is a cutting-edge analytical technique that focuses on studying the small molecules within microenvironments, up to individual cells.^28^ Improvements in instrument sensitivity and spatial resolution^29,30^ alongside the development of algorithms and software capable of effectively managing and analyzing the substantial volume of generated data^31^ have streamlined the realization of single-cell spatial metabolomics in cultured cells.^13^ The multimodal imaging method presented in this article presents a thrilling perspective on complex microenvironment spatial metabolomics. By seamlessly integrating MSI with IF on the same tissue section, and achieving a high level of spatial resolution, this approach ushers in a paradigm shift in MSI thinking. It moves away from pixel-centric considerations and, instead, centers on individual cells, providing an extraordinarily precise and detailed depiction of biological tissues. This combination delivers a comprehensive understanding of biological samples, allowing molecular features to be intricately mapped onto specific cellular structures. Such advancements are pivotal for spatial metabolomics, as they empower researchers to correlate metabolites and bioactive molecules with specific cells and tissues, unlocking new dimensions in the study of cellular dynamics.

Regarding MSI data analysis guided by IF, our work demonstrates the potential of the ROC approach, which allows for the identification of ions specific to targeted histological regions. This approach addresses the limitations of traditional univariate analysis by focusing on the discriminative ability of ions rather than average differences in ion values between groups.^19^ The “volcano ROC” plot presented in this work offers a useful way to identify quickly molecular biomarkers of IF-stained targeted cell population, like an accumulation of ceramide phosphate (**Table S2**) and SM (**figure S5**) in T cell area, a lipid family know to be very important in T cell phenotype.^32,33^ We think that the open-source Python algorithm we propose in this work might be useful for the community to achieve this goal.

In addition, another contribution of our study lies in the introduction of a non-targeted strategy with SCM metric and its combination with spatial clustering, allowing a quick overview of the different histological distributions displayed by metabolites detected by MSI. In opposition to pixel spectral clustering,^22^ this nontargeted approach offers to distinguish between ion distributions that are mere noise and those that represent genuine spatial patterns within the tissue. By quantifying the degree of spatial coherence in ion distributions, we can identify, from MSI data, regions where specific metabolites accumulate (**Figure S8d** and **f**). This measure helps investigate new spatial structures and ensures that we focus on biologically relevant ion distributions that correspond to distinct histological structures or cell populations. This computational innovation opens exciting opportunities for the discovery of previously unrecognized biomarkers and the elucidation of intricate spatial relationships in complex biological systems. Finally, the use of spatial and spectral clustering techniques provides a powerful way to highlight histological structures emerging from MSI data. These complementary approaches uncover regions of interest and complex spatial patterns, thus revealing crucial insights into the distribution of metabolites and their correlation with cellular biology.

The impact of this method on spatial metabolomics is significant because it enhances our understanding of metabolite distribution in complex and heterogeneous tissues, which is essential for studying metabolic responses in diverse biological contexts. This multimodal imaging approach and its associated computational processing have demonstrated its potential to highlight specific biological features within the spleen. For instance, it enables the identification of metabolites associated with distinct cell populations, such as T cells and MMM. ROC analysis in conjunction with IF masks revealed ions with significant and distinctive accumulation patterns within T cell-rich regions. Notably, ceramide phosphates PE-ceramides and SM accumulated prominently in T cell areas. This result sheds light on the importance of this lipid class in T cells and their potential roles in immune responses.^34,35^ Similarly, SCM and spatial clustering highlighted the accumulation of metabolites in specific but untargeted histological areas (**figure S6c**). Thanks to new IF staining and ROC analysis, we identify that some clustered ions exhibit strong associations with MMM, notably ions from spatial cluster #6, *m/z* 837.549, 863.569, and 889.581. Subsequent MS/MS imaging confirmed these ions as members of the phosphatidylinositol (PI) lipid family, respectively PI 18:0/16:0, PI 18:0/18:1, and PI 18:0/20:2 (**figure 5 and S12**). Interestingly, we observed overexpression of a lipid-containing eicosadienoic acid (EDA) in MMM. EDA is a rare fatty acid observed in animal tissues and involved in pro-inflammatory modulator production in murine macrophages.^36^ These findings offer valuable insights into the metabolic characteristics of MMM and their functional roles within the spleen.^37^ In summary, this multimodal imaging approach not only provides a comprehensive view of the spleen’s histological structures but also allows for the identification of metabolites specific to key cell populations.

In conclusion, the multimodal imaging method presented in this article opens new avenues for spatial metabolomics in MSI. It pushes the boundaries of MSI data analysis by combining chemical and histological information, using innovative algorithms, and enabling detailed cellular analysis. This advancement is critical for better comprehending the distribution of metabolites in biological tissues and its impact on cellular biology, paving the way for new discoveries in various fields of biomedical research. In addition, we provide an open-source Python library to ensure the reproducibility of our workflow. We believe that these tools will be highly valuable to the spatial omics community, particularly with the emergence of multiplexed imaging strategies such as spatial transcriptomics, cyclic immunofluorescence, and imaging mass cytometry.^38,39^ Furthermore, the algorithms we propose are not limited to MSI data (.imzML) but can also handle images in .tif format containing a large number of channels, making these tools more versatile for such multiplexed imaging applications.

## Method

### 1. Tissue sectioning

Mouse spleen biopsies were purchased from Charles Rivers (Collected from 5 mice male or female). Samples were stored at −80°C until sectioning. Sections (12 μm-thick) were cut from the spleen using an NX70 Star cryostat (Thermo Fisher Scientific) and thaw-mounted onto standard glass microscope slides.

### 2. Matrix and internal standards application

For the MALDI-MSI, a homogenous matrix layer was deposited using a home-built pneumatic sprayer^40^. Briefly, 150μL of a matrix solution was applied to the tissues under the following optimized conditions: a 0.05 mL/min matrix solution flow rate and 0.9 bar of nitrogen gas flow rate. Matrix solutions are detailed in the table below:

| Matrix | Concentration (mg/ml) | Solvent composition | Solvent Ratio | Quantity of matrix on slide (µg/mm<sup>2</sup>) |
| --- | --- | --- | --- | --- |
| DAN | 5 | Acetone/water | 7:3 | 1.5 |
| CHCA | 5 | Acetonitrile/acetone/water | 4:3:3 | 1.5 |
| DHB | 20 | Propanol/Toluene | 7:3 | 4.8 |
| 9-AA | 5 | Methanol/water | 5:5 | 1.5 |

For MS signal normalization, deuterated internal standards have been used. Matrix solutions were supplemented with 0.2 mg/ml of 1-palmitoyl-d31-2-oleoyl-sn-glycero-3-phosphate (16:0-d31-18:1 PA, Avanti Polar Lipid, [M – H]^-^, *m/z* 703.6697 ± 0.003) or 1-palmitoyl-d31-2-oleoyl-sn-glycero-3-phosphoethanolamine (16:0-d31-18:1 PE, Avanti Polar Lipid, [M - H]^-^, *m/z* 746.710 ± 0.003) for negative mode MS. A 1-palmitoyl-d31-2-oleoyl-sn-glycero-3-phosphocholine (16:0-d31-18:1 PC, Avanti Polar Lipid, *m/z* 790.7726-[M+H]^+^, *m/z* 812.754-[M+Na]^+^ and *m/z* 827.7397-[M+K]^+^) was used for positive mode MS.

### 3. MSI instrument and acquisition

Acquisitions were performed using an atmospheric pressure imaging ion source named AP-SMALDI 5 AF (TransMIT GmbH) connected to an orbital trapping mass spectrometer (QExactive Orbitrap, Thermo Fisher Scientific).^41^ This latter was operated in negative or positive ion mode at a mass resolution of 70 000 @ *m/z* 400 over a mass range of *m/z* 200-2000. The ion source is equipped with a diode laser (Flare NX343, λ = 343nm), operating at a repetition rate of 2 kHz. Imaging data were acquired in continuous mode with a pixel size of 5μm, and a speed rate of 3.7 pixels/s. Laser frequency and shots intensity were measured previously with a pyro-electrical receiver (µ-joule meter, Lasertetchnik Berlin), mounted directly on the source stage. For our acquisition parameters, source drivers adjust laser frequency to 95Hz. Concerning laser energy, 20% filter was ON and the attenuator angle was set a 39°, giving ultimately 7nJ per shot. Matrix peaks have been used as lock mass during acquisition to compensate for mass shifting.

For species MSMS identification, we define small areas in the spleen section containing splenic follicles and perform MSMS imaging by selecting parent ions and generating images of ion fragments (**Figure S12a**). Fragmentation is performed with HCD at normalized collision energy set at 28% (performed on Q-Exactive). To have a precise image of parent ions distribution, a consecutive MS1 image is performed on the same tissue section with the acquisition pattern flipped by 90° (**Figure S12b**).

### 4. Matrix rinsing, Immunofluorescence staining and acquisition

After MSI acquisition, tissue sections on glass slides are recovered from the MSI instrument and rinsed with two baths of acetone, 2 min each. Acetone rinsing was chosen over various solvent conditions according to our optimization presented in the Results section and in **Figure S2**.

Tissue sections were washed three times with Cell Stain Buffer (CSB) (Biolegend Netherlands, #420201) and incubated for 30 minutes at room temperature with Fc Block (Biolegend Netherlands, # 101320). After 3 washes with CSB buffer, tissue sections were incubated with either CD45R/B220 Alexa 488 (clone RA3-6B2, #103225), CD4 Alexa 647 (clone GK1.5 # 100424) and CD11b Brilliant Violet 421 (clone M17/0, # 101251) or CD169 BV421 (clone 3D6.112, #142421) and CD209b (clone 22D1, #53-2093-82) antibodies and CD11b Brilliant Violet 421 (clone M17/0, #101251). All antibodies were purchased from BioLegend Netherlands, except CD209b purchased from eBioscience. Fluorescently labeled antibodies were incubated overnight at 4° in CSB buffer. After three washes with CSB buffer, tissue sections were mounted with a coverslip (Zeiss #1.5H) in Prolong Glass Gold mounting media (ThermoFisher, #P36980). After immunostaining, tissue sections images were acquired in tile mode using a Zeiss LSM710 NLO or LSM900 confocal microscope equipped with a 20x Zeiss NA 0.8 objective, at a zoom of 0.6x, an image size of 512×512 pixels and with a pinhole partially opened (108.7 or 107.4μm).

### 5. Optimization of MSI workflow to turn it compatible with IF staining

First, we assessed the impact of matrix deposition and its rinsing on the IF staining. To do so, we decided to spray on tissue the four main matrices used by the MSI community on tissue section, 1,5-Diaminonaphthalene (DAN) and 9-aminoacridine (9AA) for negative mode or 2,5-Dihydroxybenzoic acid (DHB) and α-Cyano-4-hydroxycinnamic acid (CHCA) for positive mode. We then rinsed matrices after a few hours a room temperature prior to performing IF staining with antibody targeting immune cell receptors (B220/Alexa488, CD4/Alexa 647, CD11b/brilliant violet 421, **Figure S1**). For this first evaluation of matric impact, no MSI scan was performed. Interestingly, in our negative control, without any wash prior to IF staining, only CHCA did not impair the staining of all the chosen antibodies. In contrast, we see that the 3 other matrices induced strong autofluorescence in wavelength close to 400nm. We do not observe major interference for two other antibodies coupled with Alexa probes 488 and 647nm. The matrix 9AA is more difficult to remove as revealed by the strong autofluorescence in the low wavelength observed all over the slide.

We then tried to rinse MALDI matrices with different solvents (Acetone, ethanol, methanol, isopropanol, and toluene). Only acetone presented suitable results, preventing damage to the epitope of the targeted cell receptor and efficiently removing 3 matrices (9AA still present some non-specific/autofluorescent signal). All the other solvents do not remove matrices efficiently or impair the IF staining specificity. For example, the ethanol wash of DAN matrix seems acceptable for B220 and CD4 staining specific but impaired the CD11b staining.

We then optimized the MSI scan condition to limit tissue damage induced by the laser. We decided to pursue this work with the 3 matrices for which acetone wash works well: DAN, DHB, and CHCA. We scan delimited areas on spleen tissue (previously sprayed with matrices) in continuous mode at different lateral resolutions (5 and 25µm) and different laser intensities (**Figure S2**). The source driver adjusts automatically laser frequency depending on pixel rate (here 3.7pixels/s) and lateral resolution, for 5 and 25µm frequency is set at 95Hz and 430Hz, respectively. Once the different areas are scanned, the tissue is rinsed with acetone, and IF staining is performed. The laser does not touch the tissue between the different scanned areas, allowing us to properly evaluate its impact on IF staining with the different MSI scan conditions.

We observed that with the highest energy, the laser damaged the tissue and IF staining is impaired on the laser path (**Figure S2**; area #4, #5, #12, #13, and #14). The damage of the laser is even more striking at our best lateral resolution (5µm, corresponding to the size of the laser spot) where most of the IF signal is lost (**Figure S2**; area #7, #8, #17 and strong attenuation in area #16). On the other hand, we demonstrate that at lower MSI laser energy, IF signal can be preserved while getting and satisfying MSI signal (**Figure S2**; area #1, #2, #6, #9, #10, #12, #15, and #18). Interestingly CHCA matrix seems to provide the best protection from the laser, as demonstrated in areas #23 and #26 (**Figure S2**) where even at the maximum tested laser energy the IF signal is barely affected, however, the MSI signal is low. By the same time to properly evaluate the best compromise between IF staining and quality in MSI signal, we load the different MSI datasets obtained from the different areas in METASPACE2020 website to perform automated annotation. Interestingly, for all the matrices and lateral resolution conditions, we found that lower energy allows to achieve more species annotation (Table in **Figure S2**). At high laser energy, the MS signal is more intense but such as the noise, impairing proper annotation.

We observe our best annotation results with DAN in negative mode and DHB in positive mode at lower energy and for both at 25um and 5um lateral resolution. To conclude this first step of MSI optimization to be included in our multimodal workflow, we chose DAN and DHB matrices sprayed on tissue and rinsed with acetone baths. The MSI scan can be performed at any pixel size but should not exceed 7nJ.

### 6. Dataset alignment and validation

Registration was performed manually using handle transform guided by fiducial marks choosing internal standards *m/z* for the MS image (**figure S3**b). Then, the registration was assessed using the precision (p), recall (r) and F-score = 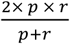 on binarized images, using an average image for the MSI dataset (**Figure S3c**).

### 7. MS data preprocessing: ion annotation, peak picking, internal standard normalization

Ion annotation was performed using METASPACE web interface^42^ based on peak exact mass and isotope profiles. MALDI images were generated using MSiReader software^43^ or Esmraldi,^18^ an in-house software; accessible at https://github.com/fgrelard/Esmraldi. Peak picking was performed on the mean spectra using the prominence of local maxima. The prominence is the height of a peak relative to the lowest contour line separating it from a higher peak within a window. This ensures peaks from signal irregularities are discarded. Here, the window size *w* depends on the chemical tolerance (*tol*, in ppm). Let *x* be the current m/z ratio, and Δ(*f*(*x*)) *= f*(*x* + *1*) *− f*(*x*) be the finite difference operator, the window size *w* is given as: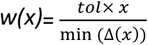. Next, we apply spectral alignment to map each pixel’s spectra onto the detected peaks. All *m/z* ratios that fall min (Δ(*x*)) within the chemical tolerance window of a detected peak are averaged and mapped to the m/z of that peak. The spectra were deisotoped. For each peak, isotopes were found by searching for peaks with a 1 Da +/-14 ppm adduct within a 8 Da window. The first peak was kept while the others were removed.

Signal normalization is performed by dividing the signal of a given ion by the signal of the deuterated internal standards. The following calculation involving MSI data used normalized data.

The resulting data structure is dense, as opposed to sparse prior to preprocessing. Thus, the images were exported in Tiff format to facilitate visualization in traditional imaging software, such as ImageJ.

### 8. IF preprocessing / Cellular region extraction

Red pulp and white pulp region are obtained by thresholding the blue channel bright field (**Figure S5a**) image in ImageJ, blue channel offering the better contrast for blood signature. Cellular regions are obtained by thresholding each channel of the IF image (**Figure S5b**) with ImageJ. The red channel corresponds to lymphocyte T, the blue channel to lymphocyte B, and the green channel to myeloid cells.

### 9. Univariate analysis

For the Volcano plots, to compare two regions of interest, pixels in those areas are considered as individual data points. For each ion, we determine the average signal and standard deviation of the signal as well as the number of pixels in selected regions. We used GraphPad Prism software to perform a multiple t.test with FDR threshold (q value) set at 1%, with Benjamini, Krieger, Yekutieli method and assuming no consistent SD. We then calculate log2(ratio) between the two regions to generate a volcano plot (**figure S4d)**.

For Receiver-Operating Characteristic (ROC) analysis^44^ of a selected MSI dataset, we considered pixels as individual data points. We generate an ROC curve for a given ion given delimited histological regions. The ROC curve is generated as follows: for each possible intensity level in the ion image, the ion image is thresholded and the specificity (Sp) and sensitivity (Se) are calculated with respect to the mask of the region. The ROC curves plot Se against 1-Sp (**Figure 2d**) from which we can determine the area under the curve (ROC AUC) which measures the specificity of an ion with regard to a region. Then, we used a routine to automate this process for all ions in all regions. ROC AUC and the ratio of ion mean intensity in the mask and its complementary are calculated, and then we plot them to obtain the “volcano-like” plot (**Figure 2f**). We consider an ion as a good biomarker of a region when its ROC AUC is above an empirical threshold of 0.8.^23^

### 10. Spatially coherent distribution filtering algorithm

At this stage, the MS image still encloses interesting spatial distributions that do not correlate with IF-targeted cells. We propose a new method that automatically identifies these ions. For all metrics, we compute binarized versions of the image over various thresholds. Indeed, the spatial distributions enclosed by each ion are apparent at different intensity levels. Intensity distributions across ions have a different number of modes, so automatic thresholding methods such as Otsu’s are not suited.^45^ In the following, we choose the set of percentiles P = [30, 50, 60, 70, 80, 90, 95]. In the following, the ion images were normalized between 0 and 255.

We define new distance-based measures. First, the **off-sample measure** (*OffM*) *e*stimates how points are spread outside of a sample that is centered in the image (**Figure S6a**). We start by estimating an off-sample image. For each ion image, the minimum distance between the center of the image and the points in the binarized image is computed. There are two distinct groups of distances with low values, corresponding to on-sample ions, and high values, corresponding to off-sample ions. We separate them using K-means with two classes. Then, a binary object *B* is built adding every point in the class of ions images with high off-sample values. Finally, the off-sample measure is defined as the average of the intensity of an ion image *I* in the binary off-sample object, *i*.*e*.,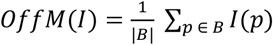. Values range from 0 to 1, with values close to 0 indicating on-sample ion images. Empirically, we select a threshold of 0.1, below which ion images are considered on-sample.

Next, our **spatially-coherent measure** (*SCM*) estimates the amount of noise with respect to spatially coherent structures (**Figure S7b**). It builds on distance transformations (DT) of the ions. The DT is a mapping of points in a binary object to their distance to the complement of the object. Let O be an object, *Ō* be its complement, the Euclidean DT maps the following value to each point 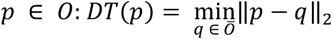 The spatially-coherent measure of an ion image estimates the prominence of noise, *i*.*e*., small structures with low DT values, with respect to larger structures. This is done by comparing the average of the DT values in the objects with the maximum of those values. It is defined as the minimum of 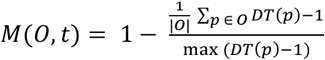 over the thersholds t ∈ P (set of percentile). Then, we range from 0 to 1 SCM values, with high values indicating spatially coherent distributions and low values corresponding to noisy ion images. We set a threshold of 0.65, above which ion images are considered to be spatially coherent.

For subsequent analysis, we used thresholded ion images. For each ion, we used as threshold t the one allowing to minimized *M*(*O,t*).

### 11. Clustering and UMAP projection

We wanted to visualize the distributions of the resulting ions and the impact of each criterion using two clustering techniques: spatial and spectral. Spatial clustering consists of grouping ion images, and spectral clustering consists of grouping pixels. Both techniques are complementary since less frequent spatial distributions might be better highlighted by spatial clustering, while small structures sharing similar spectra might be better evidenced by spectral clustering.

For spatial clustering, we used hierarchical clustering along with a projection of the clusters into a reduced space using UMAP.^25^ We used the cosine metric to build the distance matrix. We chose the number of clusters by using the inconsistency coefficient^26^ of the hierarchical clustering tree. We decide to set a threshold of 1.7 in inconsistency leading to 38 clusters. Clusters were further visualized by UMAP projection, using the previous distance matrix, modified to separate clusters in the reduced space. For each row corresponding to an ion *k*_*j*_ = *k*_*i*_ with a cluster number, the cell values associated to a cluster number *k*_*j*_*= k*_*i*_ were not modified. The other cell values (*k*_*j*_*≠ k*_*i*_) were multiplied by a factor of 10.

For spectral clustering, we used the K-means algorithm^27^ with the same number of clusters found by spatial clustering. Both clustering techniques were compared to identify the commonalities and specificities of each technique. More specifically, we estimate whether a spatial structure was found by both methods or was specific to either method. We used our distance similarity measure between the average images from the spatial clustering and the cluster images from the spectral clustering, with a threshold of 5.

### 12 Code availability

The algorithm codebase is accessible on GitHub at https://github.com/fgrelard/esmraldi. The repository contains documentation as well as examples of how to use the methods presented in this article. The methods can be found in the following files:

- Pre-processing (peak picking, spectra alignment): Esmraldi GUI, in gui/Esmraldi.pu
- ROC: examples/roc.py
- Offsample measure and SCM: examples/find_structured_images.py; and the definition of the measures in esmraldi/segmentation.py, using the *find_similar_images_distance_map* function.
- Spatial clustering with UMAP: examples/hierarchical_clustering.py

## Supporting information

Supplemental Fig 8 - Booklet

## Acknowledgments

This work was funded by a grant from the “Fondation pour la Recherche Médicale” (FRM, grant #ARF201809007123) to support L.B.’s post-doctoral fellowship. Part of this work was supported by grant #INV-004704 from the Bill and Melinda Gates Foundation to V.D. Part of this work was financially supported by the Agence National de la Recherche (France, Grants ANR-19-CE29-0010601 “MultiRaMaS”), the CNRS (International Emerging Actions (IEA) “MultIRaMaS”, and Groupement de Recherche “GdRMSI” GDR2125) to N.D. Part of this work was supported by Fondation “Toulouse Cancer Santé” (TCS, grant #2023CS130).

## Notes

https://metaspace2020.eu/project/2ff7fd6e-97fe-11ed-b433-8fd2f85a5aae?tab=datasets

https://metaspace2020.eu/project/aef7411a-97fe-11ed-b433-4f3b39d79cd5?tab=datasets

### Competing Interest Statement

The authors have declared no competing interest.

