## Supplemental Fig 8 - Booklet for "In tissue spatial single-cell metabolomics by coupling mass spectrometry imaging and immunofluorescences"

### Figure S8 : MSI dataset overview - Results of ions filtering and spatial clustering

A) IN and OFF sample peaks filtering : Montage of ion distribution images from different groups. Corresponding m/z can be read at the bottom of each tile (page 2).
B) Spatial coherence sorting : Montages of ions distribution images with high or low spatial coherence score. Corresponding m/z can be read at the bottom of each tile (page 3).
C) Montages of ion distribution images similar or different to IF masks based on ROC AUC. Corresponding m/z can be read at the bottom of each tile (page 3).
D) Average images of the different spatial clusters, associated with the corresponding list of m/z and Metaspace annotation (page 4-10).
E) KMD projection (base CH2) of relevant m/z grouped by spatial cluster (page 10).
F) Montage of m/z distributions from each spatial clusters. Corresponding m/z can be read at the bottom of each tile (page 11).

A

Off Sample ions (345)

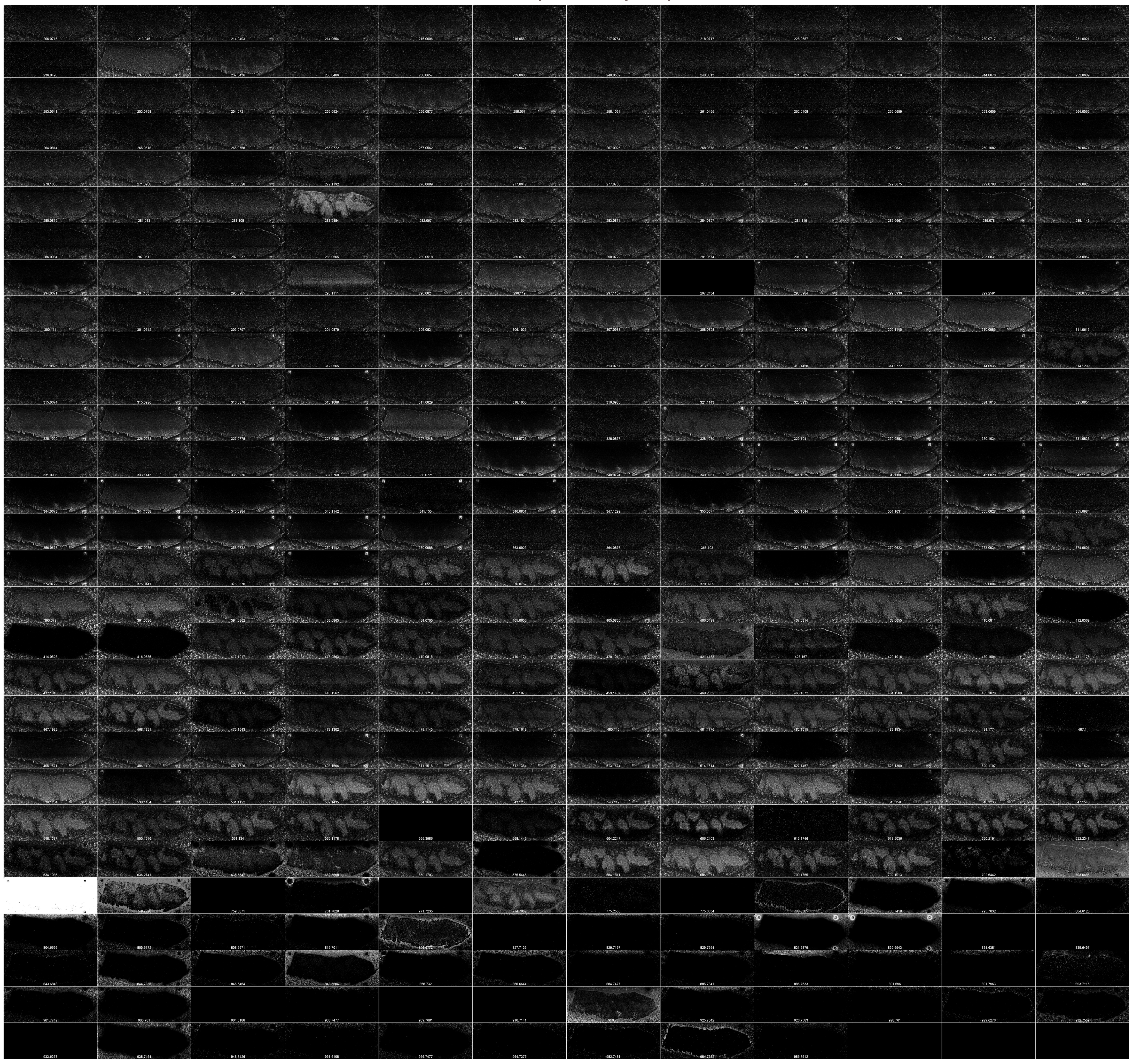

Relevant ions (224)

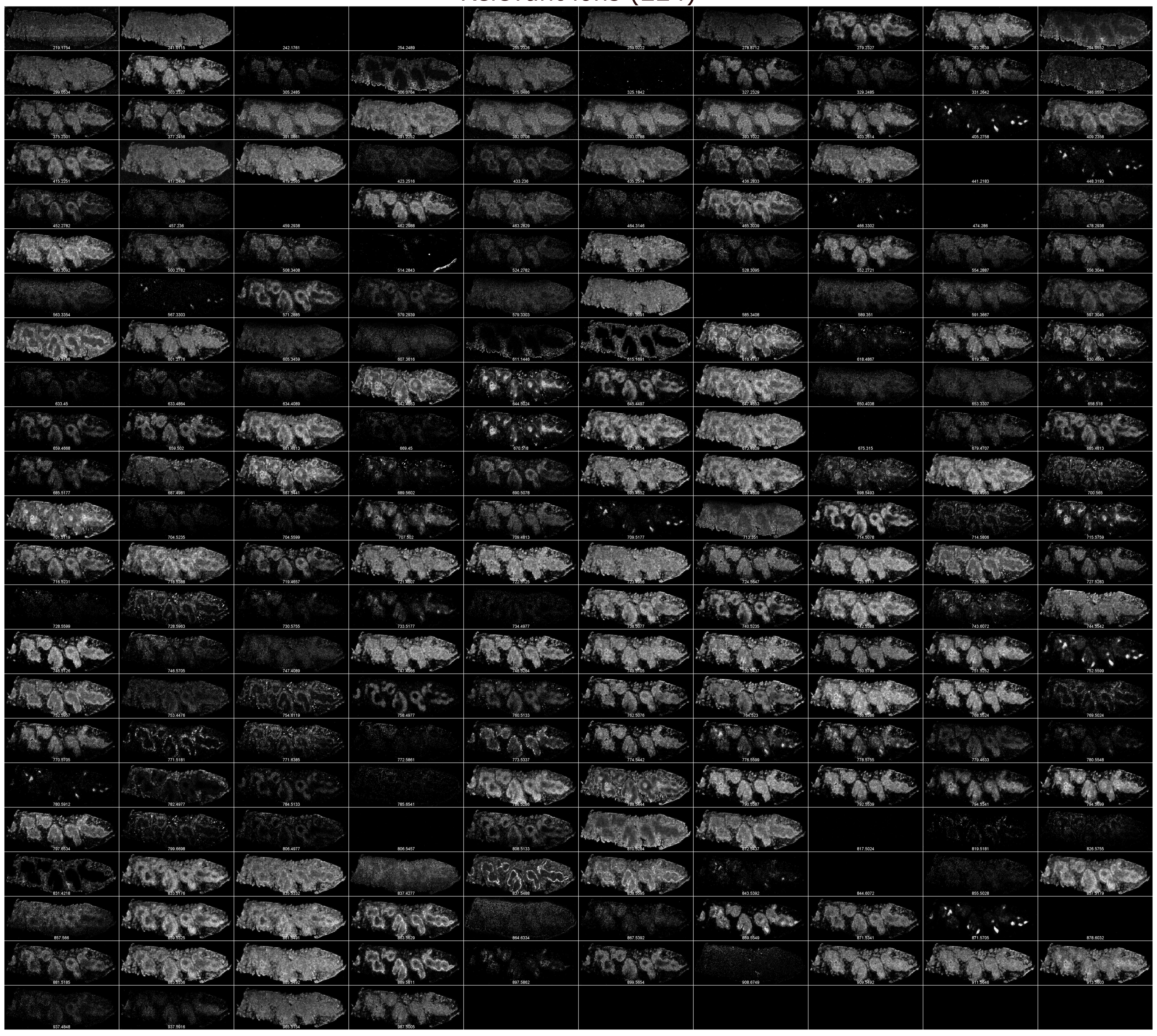

Exclude ions (15)

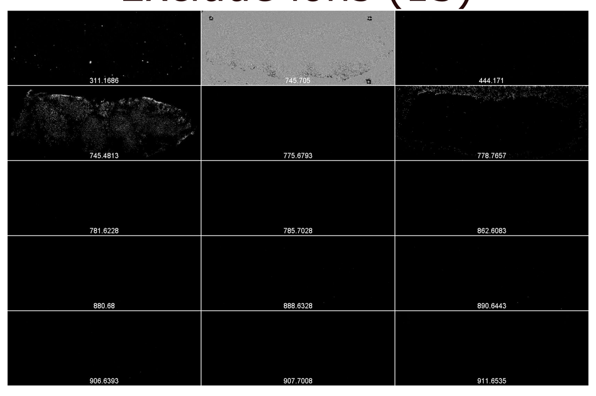

**B**

Ion distribution with high spatial coherence score (117)

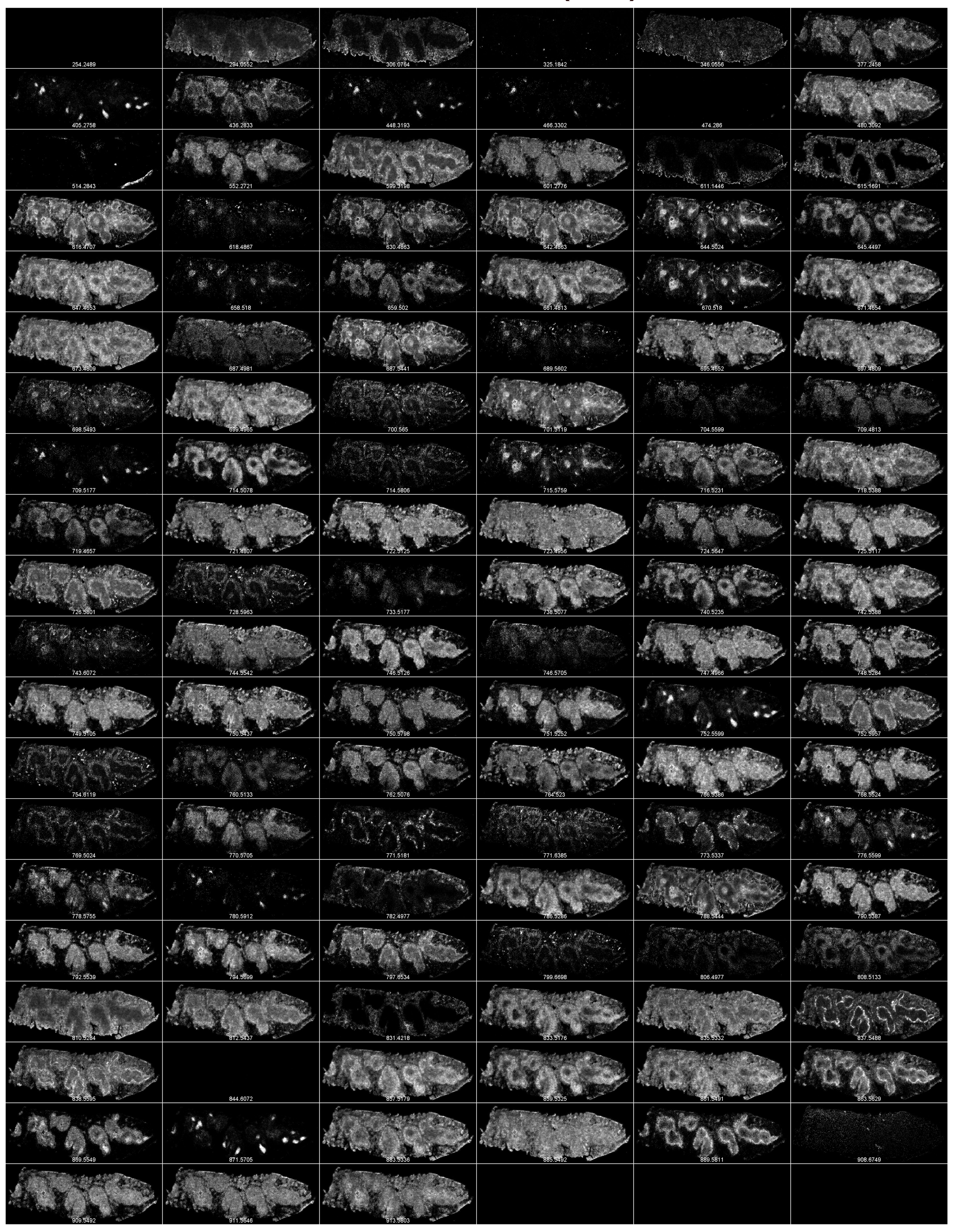

Ion distribution with low spatial coherence score (107)

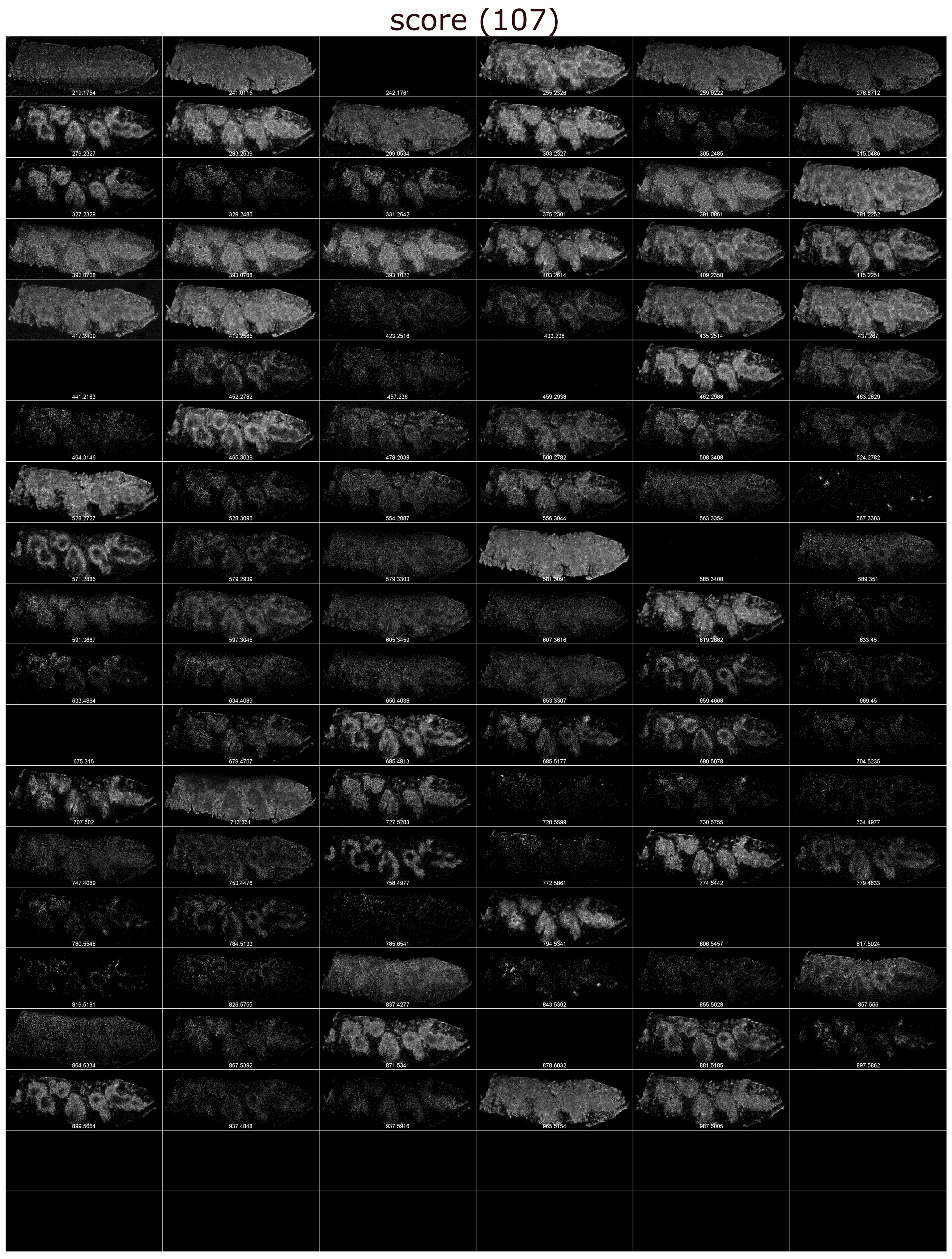

**C**

Ion distributions different to IF masks (188)

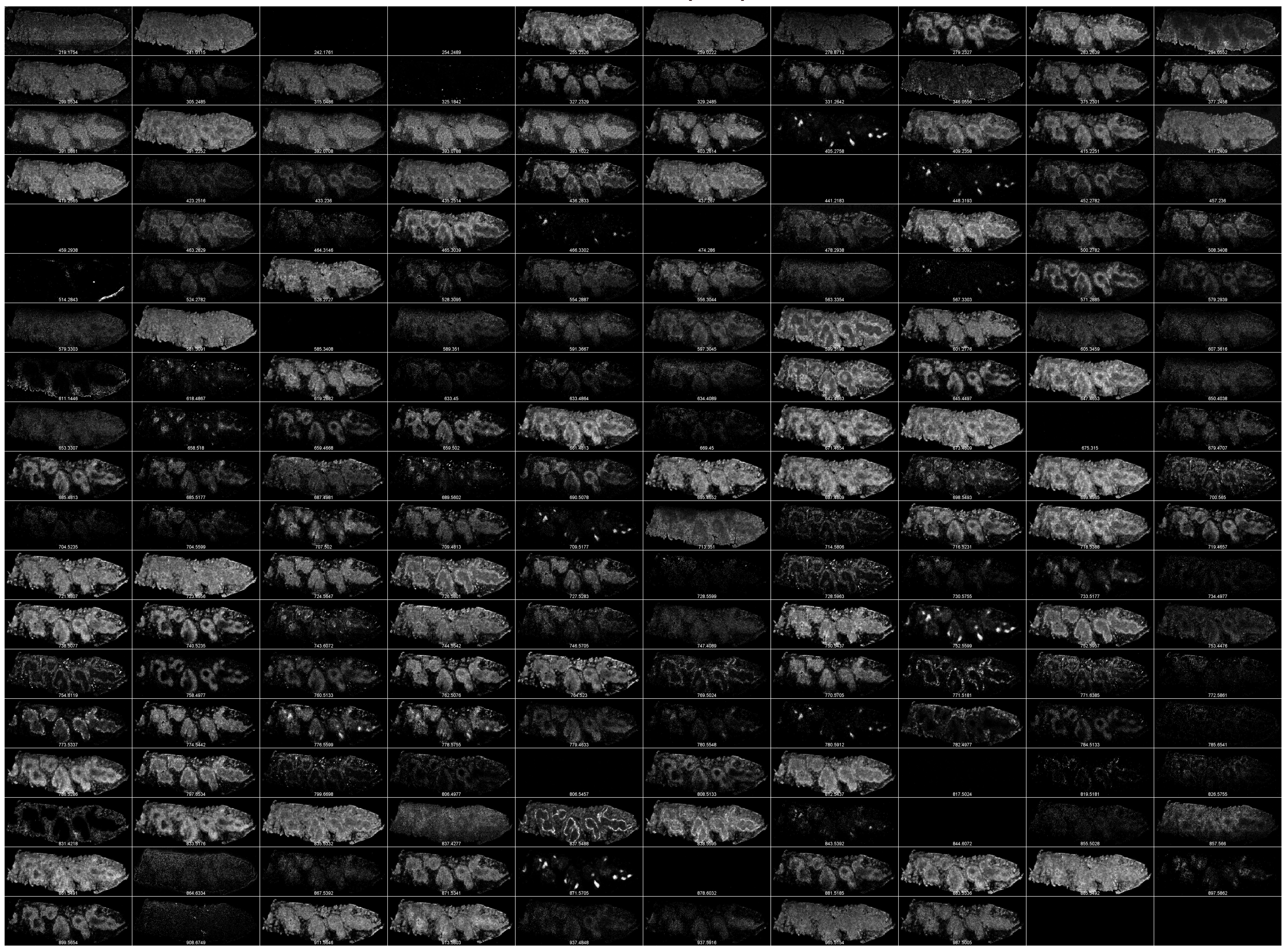

Ion distributions similar to IF masks (36)

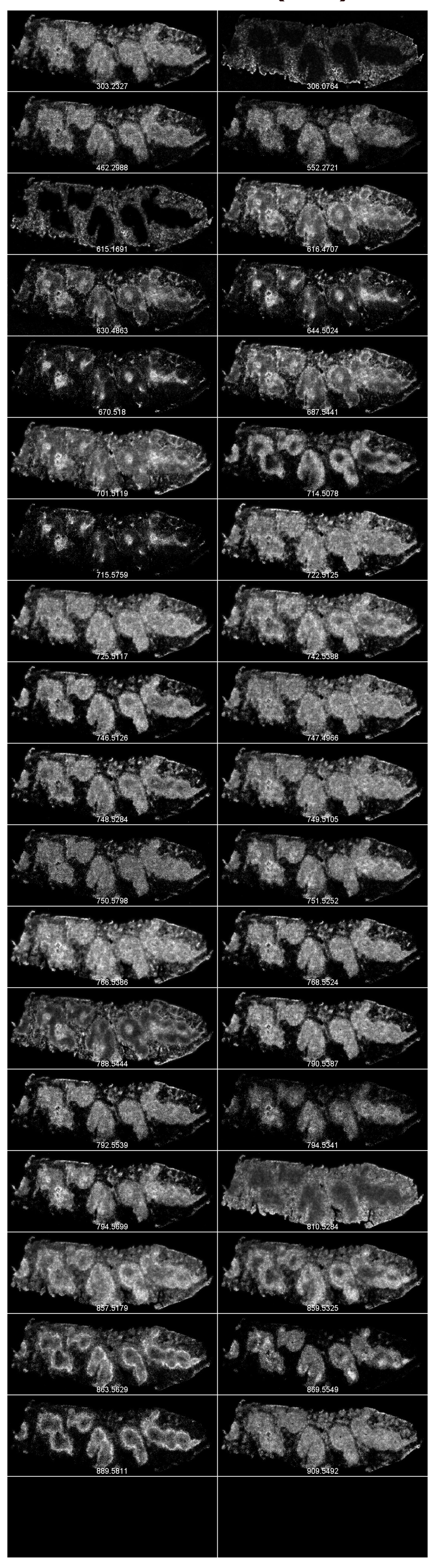

# D1

Cluster 1

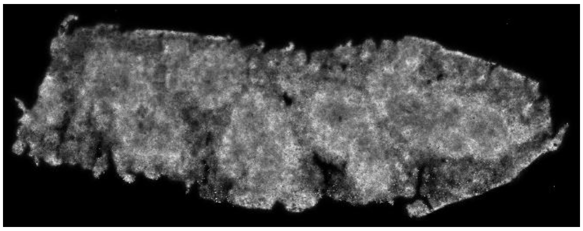

| | | | |
|---|---|---|---|
| 1 | 647.4653 | PA(32:0) | C35H69O8P-H- |
| 2 | 673.4809 | PA(34:1) | C37H71O8P-H- |
| 3 | 723.4956 | PA(38:4) | C41H73O8P-H- |
| 4 | 742.5388 | PE(36:3) | C41H78NO8P-H- |
| 5 | 766.5386 | PE(38:4) | C43H78NO8P-H- |
| 6 | 786.5286 | PS(36:2) | C42H78NO10P-H- |
| 7 | 835.5332 | PI(34:1) | C43H81O13P-H- |
| 8 | 857.5179 | PI(36:4) | C45H79O13P-H- |
| 9 | 883.5336 | PI(38:5) | C47H81O13P-H- |
| 10 | 885.5492 | PI(38:4) | C47H83O13P-H- |

Cluster 2

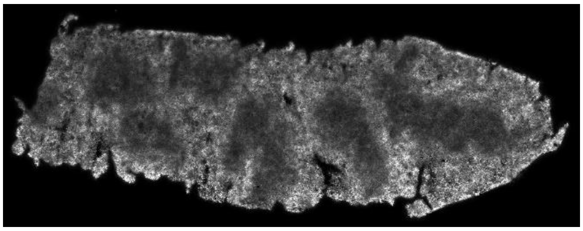

| | | | |
|---|---|---|---|
| 1 | 810.5284 | PS(38:4) | C44H78NO10P-H- |

Cluster 3

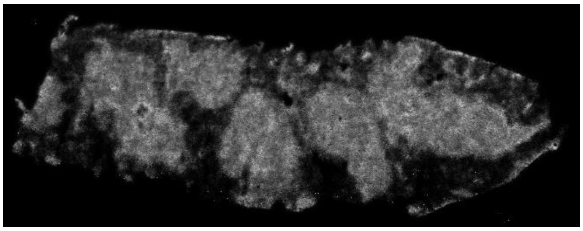

| | | | |
|---|---|---|---|
| 1 | 462.2988 | PE(P-18:1/0:0) | C23H46NO6P-H- |
| 2 | 480.3092 | PE(18:0/0:0) | C23H48NO7P-H- |
| 3 | 619.2882 | PI(20:4/0:0) | C29H49O12P-H- |
| 4 | 679.4707 | PA(P-36:4) | C39H69O7P-H- |
| 5 | 707.502 | PA(P-38:4) | C41H73O7P-H- |
| 6 | 724.5647 | 1-tetradecanyl-2-(8-[3]-ladderane-octanyl)-sn-glycerophosphocholine or ? CerP(d42:1) | C42H80NO6P-H- |
| 7 | 725.5117 | PA(38:3) | C41H75O8P-H- |
| 8 | 727.5283 | PA(38:2) | C41H77O8P-H- |
| 9 | 747.4966 | PA(40:6) | C43H73O8P-H- |
| 10 | 749.5105 | PA(40:5) | C43H75O8P-H- |
| 11 | 750.5798 | ?CerP(d44:0) or PC(P-18:0/P-18:1)? | |
| 12 | 751.5252 | PA(40:4) | C43H77O8P-H- |
| 13 | 913.5803 | PI(40:4) | C49H87O13P-H- |

Cluster 4

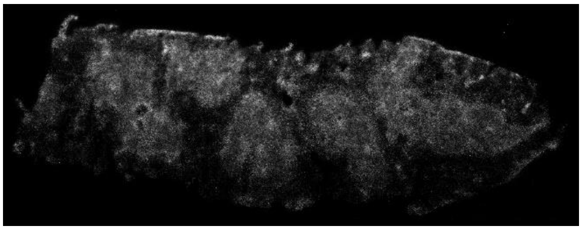

| | | | |
|---|---|---|---|
| 1 | 508.3408 | PE(20:0/0:0) | C25H52NO7P-H- |
| 2 | 743.6072 | SM(d18:1/19:0) | C42H85N2O6P-H- |
| 3 | 744.5542 | PE(36:1) | C41H80NO8P-H- |
| 4 | 746.5705 | PE(36:0) | C41H82NO8P-H- |
| 5 | 770.5705 | PE(38:2) | C43H82NO8P-H- |

Cluster 5

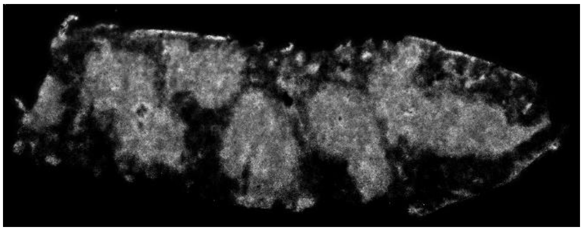

| | | | |
|---|---|---|---|
| 1 | 722.5125 | ? | |
| 2 | 750.5437 | PE(P-38:4) | C43H78NO7P-H- |
| 3 | 768.5524 | PE(38:3) | C43H80NO8P-H- |
| 4 | 774.5442 | PE(P-40:6) | C45H78NO7P-H- |
| 5 | 776.5599 | PE(P-40:5) | C45H80NO7P-H- |
| 6 | 778.5755 | PE(P-40:4) | C45H82NO7P-H- |
| 7 | 790.5387 | PE(40:6) | C45H78NO8P-H- |
| 8 | 792.5539 | PE(40:5) | C45H80NO8P-H- |
| 9 | 794.5699 | PE(40:4) | C45H82NO8P-H- |

Cluster 6

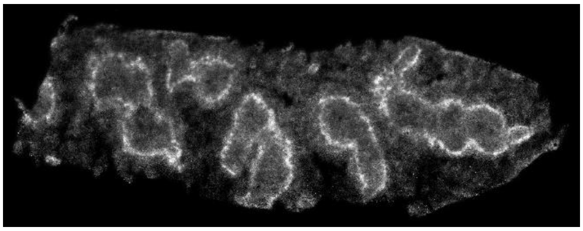

| | | | |
|---|---|---|---|
| 1 | 837.5488 | PI(34:0) | C43H83O13P-H- |
| 2 | 838.5595 | PS(40:4) | C46H82NO10P-H- |
| 3 | 863.5629 | PI(36:1) | C45H85O13P-H- |
| 4 | 889.5811 | PI(38:2) | C47H87O13P-H- |

D2

Cluster 7

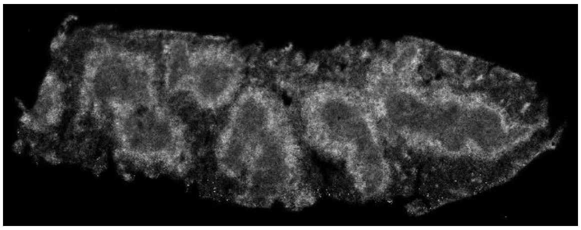

| | | | |
|---|---|---|---|
| 1 | 436.2833 | PE(P-16:0/0:0) | C21H44NO6P-H- |
| 2 | 465.3039 | Cholesterol sulfate | C27H46O4S-H- |
| 3 | 571.2885 | PI(16:0/0:0) | C25H49O12P-H- |
| 4 | 599.3198 | PI(18:0/0:0) | C27H53O12P-H- |
| 5 | 726.5801 | CerP(d42:2) | C42H82NO6P-H- |
| 6 | 752.5957 | PC(P-18:1/P-18:1) or ?CerP(44:1) | C44H84NO6P-H- |

Cluster 8

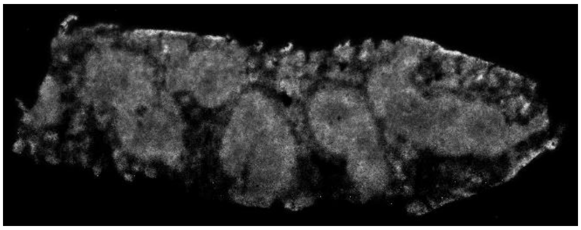

| | | | |
|---|---|---|---|
| 1 | 687.4981 | DG(42:10) or PA(35:1) | C45H68O5-H- |
| 2 | 695.4652 | PA(36:4) | C39H69O8P-H- |
| 3 | 709.4813 | PA(37:4) | C40H71O8P-H- |
| 4 | 716.5231 | PE(34:1) | C39H76NO8P-H- |
| 5 | 719.4657 | PA(36:6) | C41H69O8P-H- |
| 6 | 721.4807 | PA(38:5) | C41H71O8P-H- |
| 7 | 762.5076 | PE(38:6) | C43H74NO8P-H- |
| 8 | 764.523 | PE(38:5) | C43H76NO8P-H- |
| 9 | 808.5133 | PS(38:5) | C44H76NO10P-H- |
| 10 | 911.5646 | PI(40:5) | C49H85O13P-H- |

Cluster 9

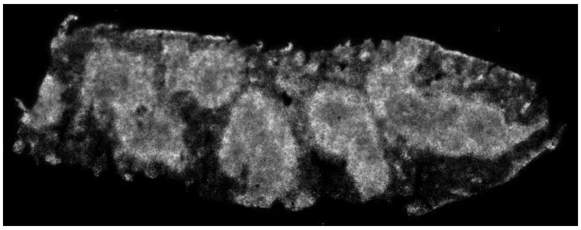

| | | | |
|---|---|---|---|
| 1 | 718.5388 | PE(34:0) | C39H78NO8P-H- |
| 2 | 738.5077 | PE(36:4) | C41H74NO8P-H- |
| 3 | 746.5126 | PE(P-38:6) | C43H74NO7P-H- |
| 4 | 748.5284 | ?PE(P-38:5) | |
| 5 | 797.6534 | SM(d41:2) | C46H91N2O6P-H- |
| 6 | 812.5437 | PS(38:3) | C44H80NO10P-H- |

Cluster 10

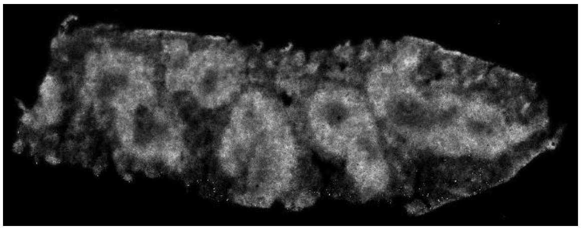

| | | | |
|---|---|---|---|
| 1 | 645.4497 | PA(32:1) | C35H67O8P-H- |
| 2 | 661.4813 | PA(33:0) or DG (40:9) | C36H71O8P-H- |
| 3 | 671.4654 | PA(34:2) | C37H69O8P-H- |
| 4 | 685.4813 | PA(35:2) | C38H71O8P-H- |
| 5 | 697.4809 | PA(36:3) | C39H71O8P-H- |
| 6 | 699.4965 | PA(36:2) | C39H73O8P-H- |
| 7 | 714.5078 | PE(34:2) | C39H74NO8P-H- |
| 8 | 740.5235 | PE(36:3) | C41H76NO8P-H- |
| 9 | 833.5176 | ?PI(34:2) | |
| 10 | 859.5325 | PI(36:3) | C45H81O13P-H- |
| 11 | 861.5491 | PI(36:2) | C45H83O13P-H- |
| 12 | 871.5341 | PI(37:4) | C46H81O13P-H- |
| 13 | 881.5185 | PI(38:6) | C47H79O13P-H- |
| 14 | 899.5654 | PI(39:4) | C48H85O13P-H- |

Cluster 11

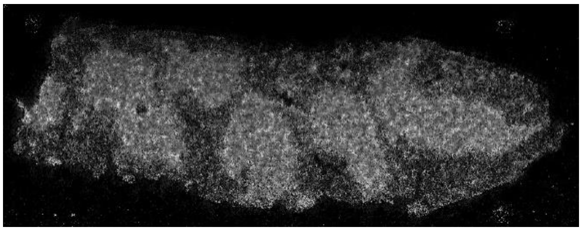

| | | | |
|---|---|---|---|
| 1 | 391.0861 | {4-[(E)-2-{3,5-dihydroxy-4-[(1E)-3-hydroxy-3-methylbut-1-en-1-yl]phenyl}ethenyl]phenyl}oxidanesulfonic acid | C19H20O7S-H- |
| 2 | 392.0708 | ? | |
| 3 | 393.0788 | ? | |
| 4 | 393.1022 | ? | |

Cluster 12

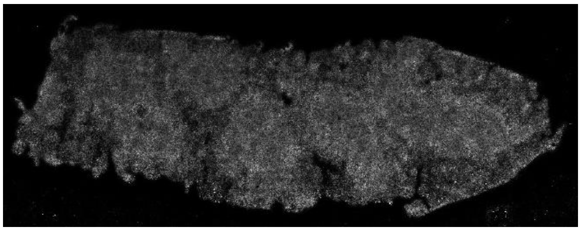

| | | | |
|---|---|---|---|
| 1 | 241.0115 | Inositol cyclic phosphate | C6H11O8P-H- |
| 2 | 259.0222 | Galactose 1-phosphate | C6H13O9P-H- |
| 3 | 299.0534 | ? | |
| 4 | 315.0486 | ? | |
| 5 | 375.2301 | ? | |

# D3

### Cluster 13

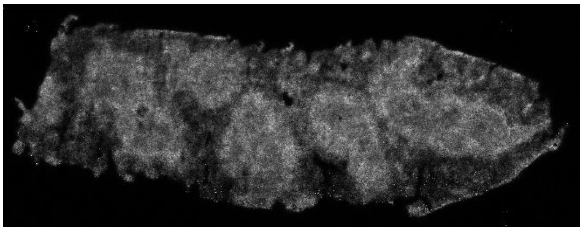

| | | | |
|---|---|---|---|
| 1 | 255.2326 | Palmitic acid 16:0 | C16H32O2-H- |
| 2 | 279.2327 | Linoleic acid 18:2 | C18H32O2-H- |
| 3 | 283.2639 | Stearic acid 18:0 | C18H36O2-H- |
| 4 | 303.2327 | Arachidonic acid 20:4 | C20H32O2-H- |
| 5 | 327.2329 | 22:6 | C22H32O2-H- |
| 6 | 331.2642 | 22:4 | C22H36O2-H- |
| 7 | 391.2252 | CPA(16:0/0:0) | C19H37O6P-H- |
| 8 | 409.2358 | LysoPA(0:0/16:0) | C19H39O7P-H- |
| 9 | 415.2251 | CPA(18:2/0:0) | C21H37O6P-H- |
| 10 | 417.2409 | CPA(18:1/0:0) | C21H39O6P-H- |
| 11 | 419.2565 | CPA(18:0/0:0) | C21H41O6P-H- |
| 12 | 435.2514 | PA(0:0/18:1) | C21H41O7P-H- |
| 13 | 437.267 | LPA(0:0/18:0) | C21H43O7P-H- |

### Cluster 14

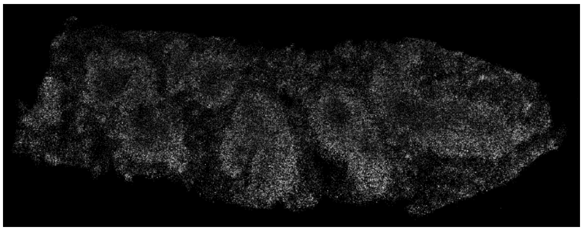

| | | | |
|---|---|---|---|
| 1 | 278.8712 | ? | |
| 2 | 753.4476 | PPA(34:1) | C37H72O11P2-H- |
| 3 | 779.4633 | PPA(36:2) | C39H74O11P2-H- |
| 4 | 937.4848 | PIP(36:4) | C45H80O16P2-H- |

### Cluster 15

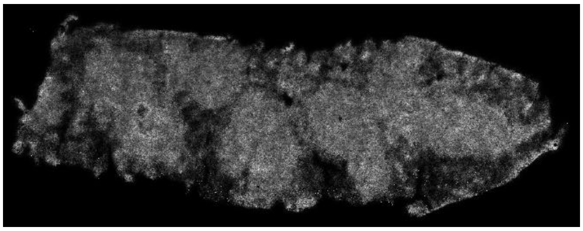

| | | | |
|---|---|---|---|
| 1 | 403.2614 | ? | |
| 2 | 581.3091 | PI(P-18:1/0:0) | C27H51O11P-H- |
| 3 | 601.2776 | ? | |
| 4 | 909.5492 | PI(40:6) | C49H83O13P-H- |
| 5 | 965.5154 | PIP(38:4) | C47H84O16P2-H- |
| 6 | 987.5005 | PIP(40:7) | |

### Cluster 16

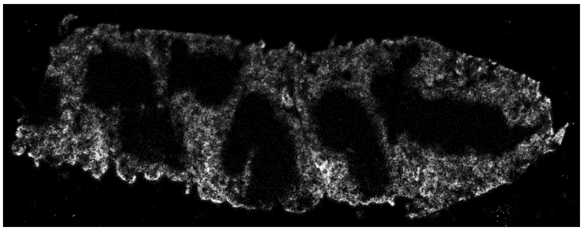

| | | | |
|---|---|---|---|
| 1 | 306.0764 | Glutathione | C10H17N3O6S-H- |
| 2 | 611.1446 | Oxidized glutathione | C20H32N6O12S2-H- |
| 3 | 615.1691 | $^{56}$Fe-HEME | |
| 4 | 831.4218 | PGP(35:6) | C41H70O13P2-H- |

### Cluster 17

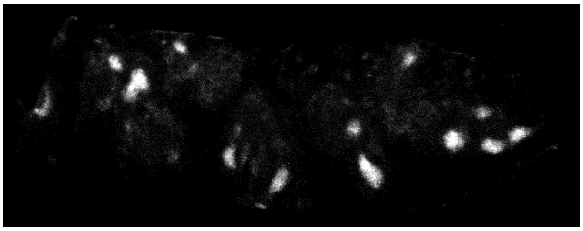

| | | | |
|---|---|---|---|
| 1 | 405.2758 | N-linolenoyl-glutamine | C23H38N2O4-H- |
| 2 | 448.3193 | 3alpha,7alpha,12alpha-trihydroxy-5beta-cholestanate | C27H45O5-H- |
| 3 | 709.5177 | PA(P-38:3) | C41H75O7P-H- |
| 4 | 752.5599 | PE(P-38:3) | C43H80NO7P-H- |
| 5 | 780.5912 | PC(P-40:3) | C45H84NO7P-H- |
| 6 | 871.5705 | PI(P-38:3) | C47H85O12P-H- |

### Cluster 18

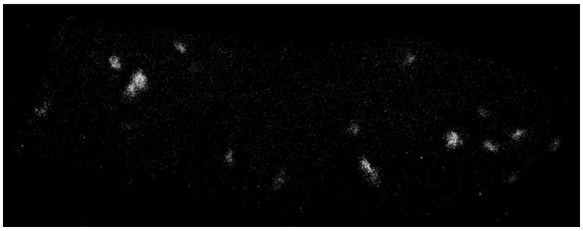

| | | | |
|---|---|---|---|
| 1 | 242.1761 | N-Undecanoylglycine | C13H25NO3-H- |
| 2 | 254.2489 | Palmitamide | C16H33NO-H- |
| 3 | 441.2183 | Rhodamine B | C28H30N2O3-H- |
| 4 | 459.2938 | (6R)-vitamin D2 6,19-sulfur dioxide adduct / (6R)-ergocalciferol 6,19-sulfur dioxide adduct | C28H44O3S-H- |
| 5 | 466.3302 | PE(O-18:0/0:0) | C23H50NO6P-H- |
| 6 | 474.286 | Hydrocortamate | C27H41NO6-H- |
| 7 | 567.3303 | ? | |
| 8 | 585.3408 | PI(O-18:0/0:0) | C27H55O11P-H- |
| 9 | 675.315 | PI(27:4) | C32H53O13P-H- |
| 10 | 806.5457 | (3'-sulfo)Galbeta-Cer(d18:1/18:0) | C42H81NO11S-H- |
| 11 | 817.5024 | PG(40:8) | C46H75O10P-H- |
| 12 | 844.6072 | PS(40:1) | C46H88NO10P-H- |
| 13 | 878.6032 | (3'-sulfo)Galbeta-Cer(d18:1/22:0(2OH)) | C46H89NO12S-H- |

# D4

### Cluster 19

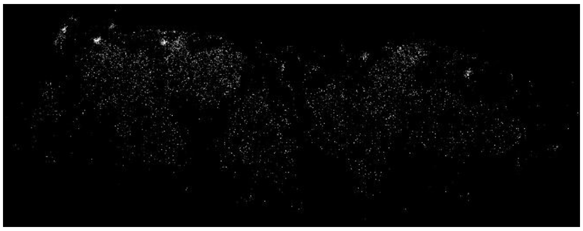

| | | | |
|---|---|---|---|
| 1 | 728.5599 | PE(P-36:1) | C41H80NO7P-H- |

### Cluster 20

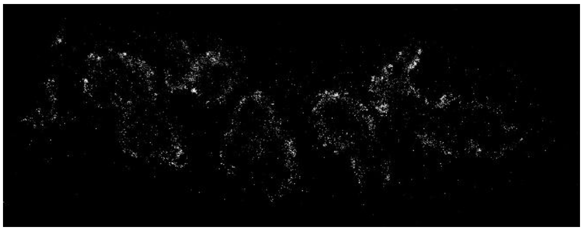

| | | | |
|---|---|---|---|
| 1 | 819.5181 | PG(40:7) | C46H77O10P-H- |

### Cluster 21

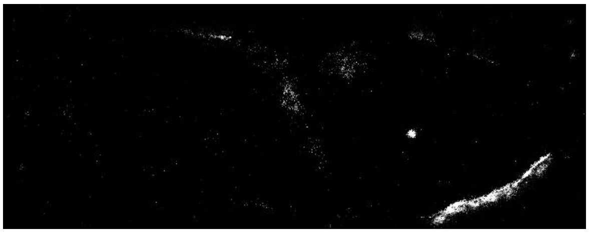

| | | | |
|---|---|---|---|
| 1 | 514.2843 | Taurocholic acid | C26H45NO7S-H- |

### Cluster 22

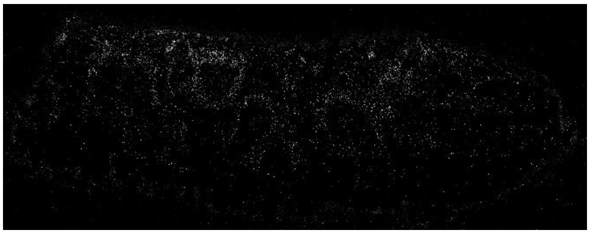

| | | | |
|---|---|---|---|
| 1 | 785.6541 | SM(d40:1) | C45H91N2O6P-H- |

### Cluster 23

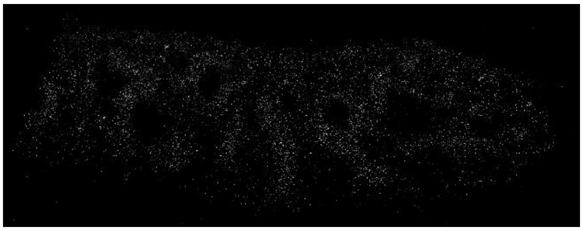

| | | | |
|---|---|---|---|
| 1 | 734.4977 | PS(32:0) | C38H74NO10P-H- |

### Cluster 24

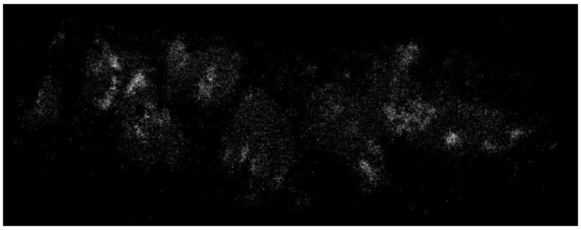

| | | | |
|---|---|---|---|
| 1 | 843.5392 | PI(P-36:3) | C45H81O12P-H- |
| 2 | 897.5862 | PI(P-40:5) | C49H87O12P-H- |

D5

Cluster 25

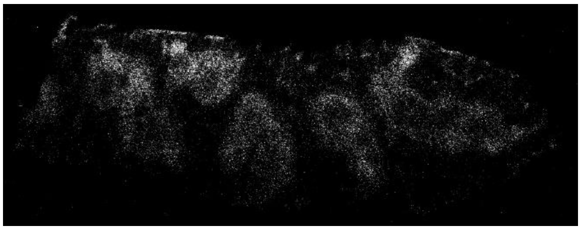

| | | | |
|---|---|---|---|
| 1 | 633.45 | PA(31:0) | C34H67O8P-H- |
| 2 | 669.45 | PA(34:3) | C37H67O8P-H- |
| 3 | 704.5235 | PE(33:0) | C38H76NO8P-H- |
| 4 | 730.5755 | PE(P-36:0) | C41H82NO7P-H- |
| 5 | 772.5861 | PE(38:1) | C43H84NO8P-H- |
| 6 | 780.5548 | PE(39:4) | C44H80NO8P-H- |

Cluster 26

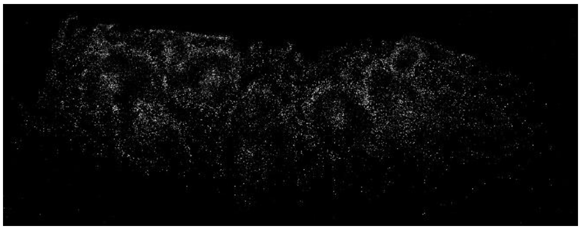

| | | | |
|---|---|---|---|
| 1 | 826.5755 | PE(P-44:8) | C49H82NO7P-H- |

Cluster 27

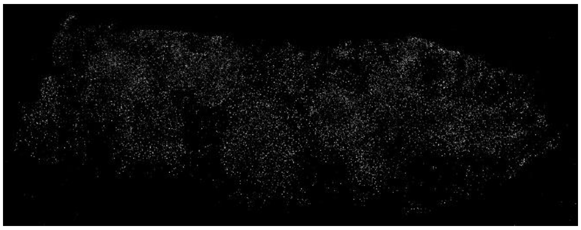

| | | | |
|---|---|---|---|
| 1 | 855.5028 | PI(36:5) | C45H77O13P-H- |

Cluster 28

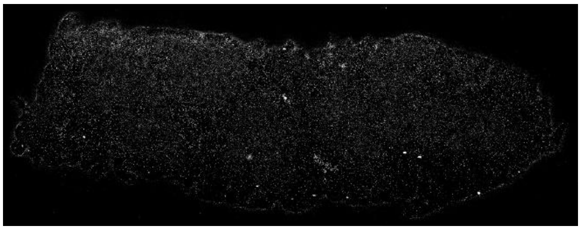

| | | | |
|---|---|---|---|
| 1 | 325.1842 | 2-Dodecylbenzenesulfonic acid | C18H30O3S-H- |
| 2 | 864.6334 | PI-Cer(d40:0) | C46H92NO11P-H- |
| 3 | 908.6749 | PS(P-46:3) | C52H96NO9P-H- |

Cluster 29

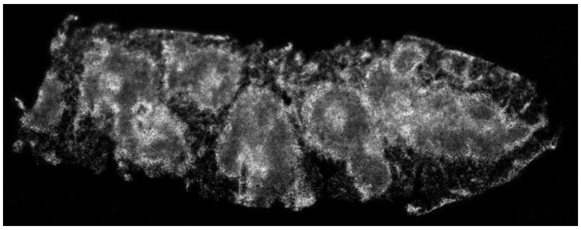

| | | | |
|---|---|---|---|
| 1 | 616.4707 | CerP(d34:1) | C34H68NO6P-H- |
| 2 | 630.4863 | CerP(d35:1) | C35H70NO6P-H- |
| 3 | 642.4863 | CerP(d36:2) | C36H70NO6P-H- |
| 4 | 687.5441 | PE-Cer(d36:1) | C38H77N2O6P-H- |

Cluster 30

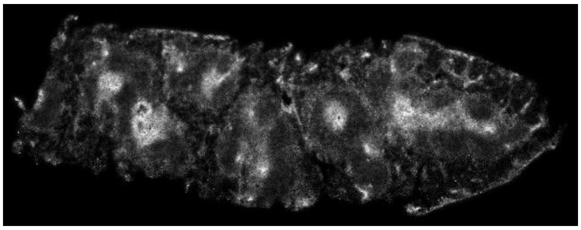

| | | | |
|---|---|---|---|
| 1 | 644.5024 | CerP(d36:1) | C36H72NO6P-H- |
| 2 | 670.518 | CerP(d38:2) | C38H74NO6P-H- |
| 3 | 701.5119 | PA(38:1) | C39H75O8P-H- |
| 4 | 715.5759 | PE-Cer(d38:1) | C40H81N2O6P-H- |
| 5 | 788.5444 | PS(36:1) | C42H80NO10P-H- |

# D6

### Cluster 31

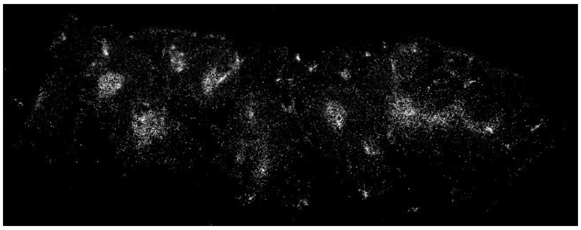

| | | | |
|---|---|---|---|
| 1 | 658.518 | CerP(d37:1) | C37H74NO6P-H- |

### Cluster 32

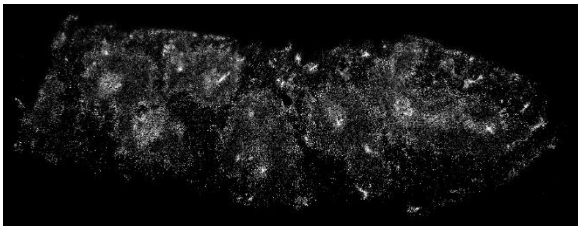

| | | | |
|---|---|---|---|
| 1 | 698.5493 | CerP(d40:2) | C40H78NO6P-H- |

### Cluster 33

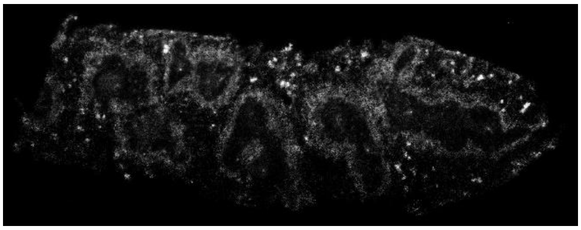

| | | | |
|---|---|---|---|
| 1 | 700.565 | CerP(d40:1) | C40H80NO6P-H- |
| 2 | 714.5806 | ?CerP(d41:1) | |
| 3 | 728.5963 | CerP(d42:1) | C42H84NO6P-H- |
| 4 | 754.6119 | CerP(d44:2) | C44H86NO6P-H- |
| 5 | 771.6385 | PE-Cer(d42:1) | C44H89N2O6P-H- |
| 6 | 799.6698 | PE-Cer(d44:1) | C46H93N2O6P-H- |

### Cluster 34

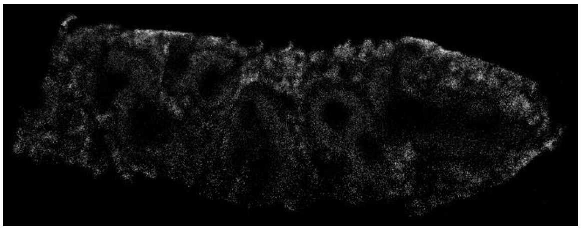

| | | | |
|---|---|---|---|
| 1 | 782.4977 | PS(36:4) | C42H74NO10P-H- |
| 2 | 806.4977 | PS(38:6) | C44H74NO10P-H- |

### Cluster 35

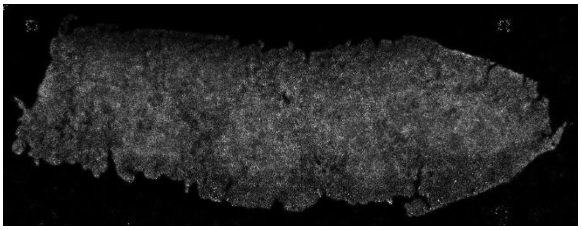

| | | | |
|---|---|---|---|
| 1 | 219.1754 | 2-trans,6-trans-Farnesal | C15H24O-H- |
| 2 | 294.0552 | ? | |
| 3 | 346.0556 | Adenosine monophosphate | C10H14N5O7P-H- |
| 4 | 563.3354 | PG(22:2/0:0) | C28H53O9P-H- |
| 5 | 579.3303 | ? | |
| 6 | 589.351 | PG(P-24:2) | C30H55O9P-H- |
| 7 | 591.3667 | PG(P-24:1) | C30H57O9P-H- |
| 8 | 605.3459 | PG(24:2) | C30H55O10P-H- |
| 9 | 607.3616 | ? PG(24:1) | |
| 10 | 650.4038 | PS(26:0) | C32H62NO10P-H- |
| 11 | 653.3307 | PI(21:1) | C30H55O13P-H- |
| 12 | 713.351 | ? | |
| 13 | 747.4089 | PI(28:3) | C37H65O13P-H- |
| 14 | 937.5916 | PGP(42:2) | C51H86O15-H- |

### Cluster 36

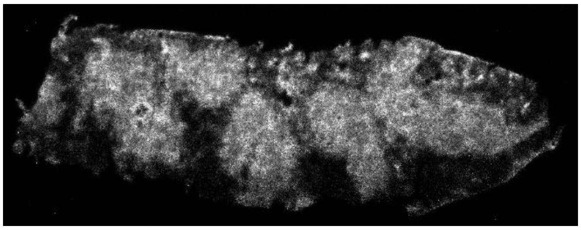

| | | | |
|---|---|---|---|
| 1 | 457.236 | 1-(5Z,8Z,11Z,14Z-eicosatetraenoyl)-sn-glycero-3-phosphate | C23H39O7P-H- |
| 2 | 464.3146 | PE(P-18:0) | C23H48NO6P-H- |
| 3 | 478.2938 | LysoPE(0:0/18:1) | C23H46NO7P-H- |
| 4 | 500.2782 | LysoPE(0:0/20:4) | C25H44NO7P-H- |
| 5 | 524.2782 | LysoPE(0:0/22:6) | C27H44NO7P-H- |
| 6 | 528.2727 | PE(21:4) | C26H44NO8P-H- |
| 7 | 528.3095 | LysoPE(0:0/22:4) | C27H48NO7P-H- |
| 8 | 552.2721 | Vignatic acid A | C30H39N3O7-H- |
| 9 | 554.2887 | PE(23:5) | C28H46NO8P-H- |
| 10 | 556.3044 | PE(23:4) | C28H48NO8P-H- |
| 11 | 618.4867 | PC(P-26:0/0:0) or CerP(d18:0/16:0) | C34H70NO6P-H- |
| 12 | 634.4089 | PS(P-26:0) | C32H62NO9P-H- |
| 13 | 689.5602 | PE-Cer(d36:0) | C38H79N2O6P-H- |
| 14 | 794.5341 | PS(P-38:4) | C44H78NO9P-H- |
| 15 | 837.4277 | Talinumoside I | C43H66O16-H- |

D7

Cluster 37

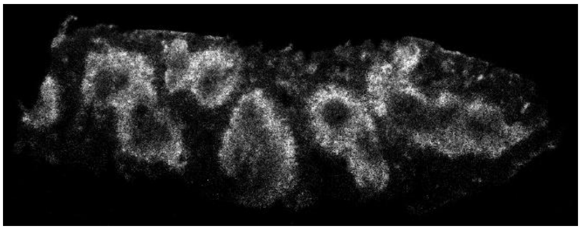

| | | | |
|---|---|---|---|
| 1 | 452.2782 | LysoPE(0:0/16:0) | C21H44NO7P-H- |
| 2 | 633.4864 | PA(O-32:0) | C35H71O7P-H- |
| 3 | 659.4668 | DG(40:10) or PA(33:1) | C43H64O5-H- |
| 4 | 685.5177 | PA(P-36:1) | C39H75O7P-H- |
| 5 | 690.5078 | PE(32:0) | C37H74NO8P-H- |
| 6 | 704.5599 | PE(O-34:0) | C39H80NO7P-H- |
| 7 | 758.4977 | PS(34:2) | C40H74NO10P-H- |
| 8 | 760.5133 | PS(34:1) | C40H76NO10P-H- |
| 9 | 769.5024 | PG(36:4) | C42H75O10P-H- |
| 10 | 771.5181 | PG(36:3) | C42H77O10P-H- |
| 11 | 773.5337 | PG(36:2) | C42H79O10P-H- |
| 12 | 784.5133 | PS(36:3) | C42H76NO10P-H- |
| 13 | 857.566 | di-trans,poly-cis-decaprenyl diphosphate or PG(P-44:8) | C50H84O7P2-H- |

Cluster 38

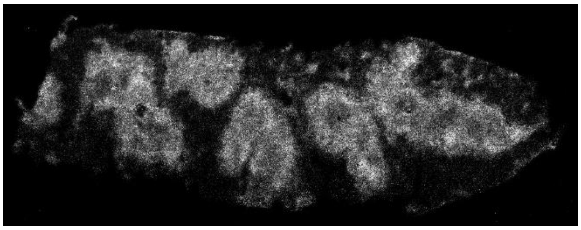

| | | | |
|---|---|---|---|
| 1 | 305.2485 | Eicosatrienoic acid 20:3 | C20H34O2-H- |
| 2 | 329.2485 | 22:5 | C22H34O2-H- |
| 3 | 377.2458 | ? PA(O-16:0/0:0) –$H_2O$ | |
| 4 | 423.2516 | PA(17:0/0:0) | C20H41O7P-H- |
| 5 | 433.236 | PA(18:2/0:0) | C21H39O7P-H- |
| 6 | 463.2829 | PA(20:1/0:0) | C23H45O7P-H- |
| 7 | 579.2939 | ? LysoPI(18:1/0:0) – $H_2O$ | |
| 8 | 597.3045 | PI(18:1/0:0) | C27H51O12P-H- |
| 9 | 659.502 | PA(P-34:0) | C37H73O7P-H- |
| 10 | 733.5177 | PA(P-40:6) | C43H75O7P-H- |
| 11 | 867.5392 | PI(P-38:5) | C47H81O12P-H- |
| 12 | 869.5549 | PI(P-38:4) | C47H83O12P-H- |

E

KMD projection

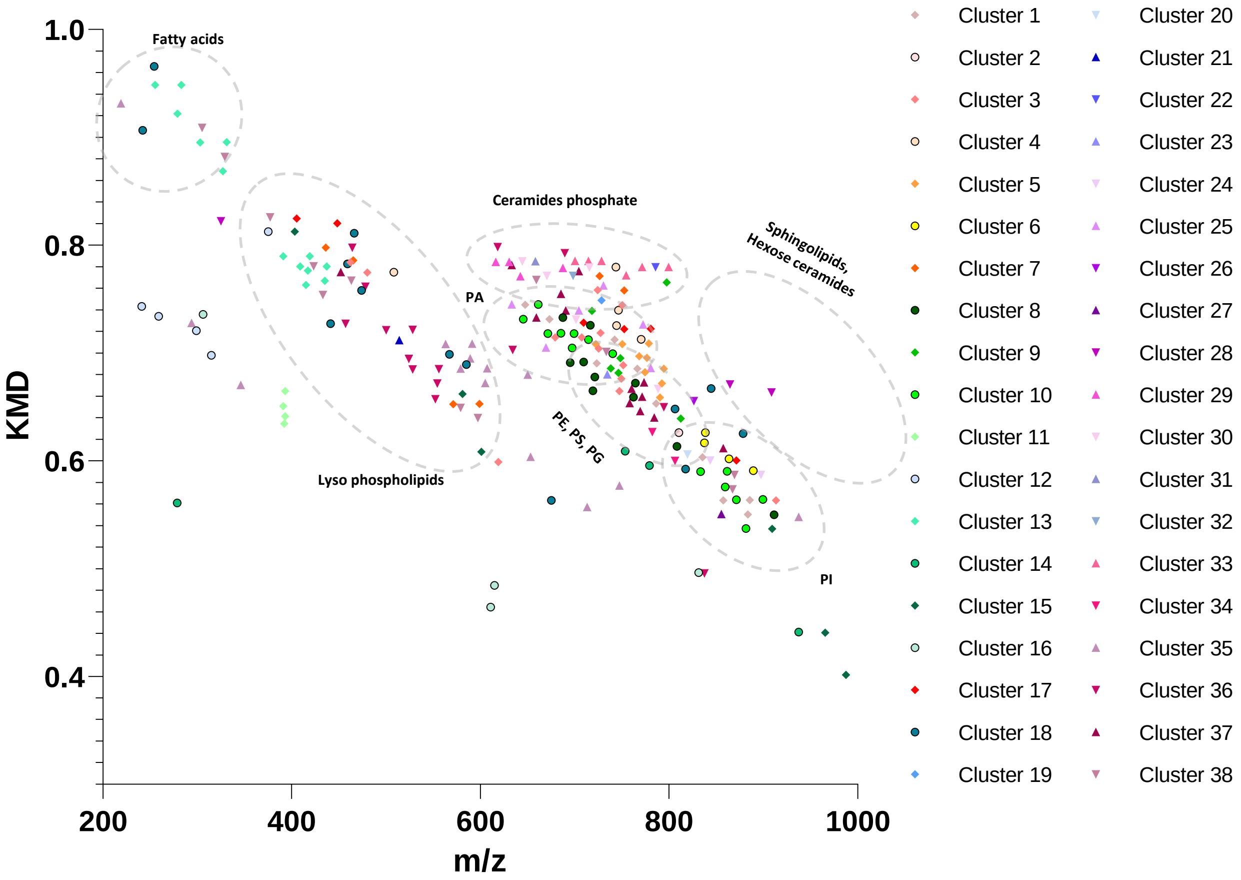

F

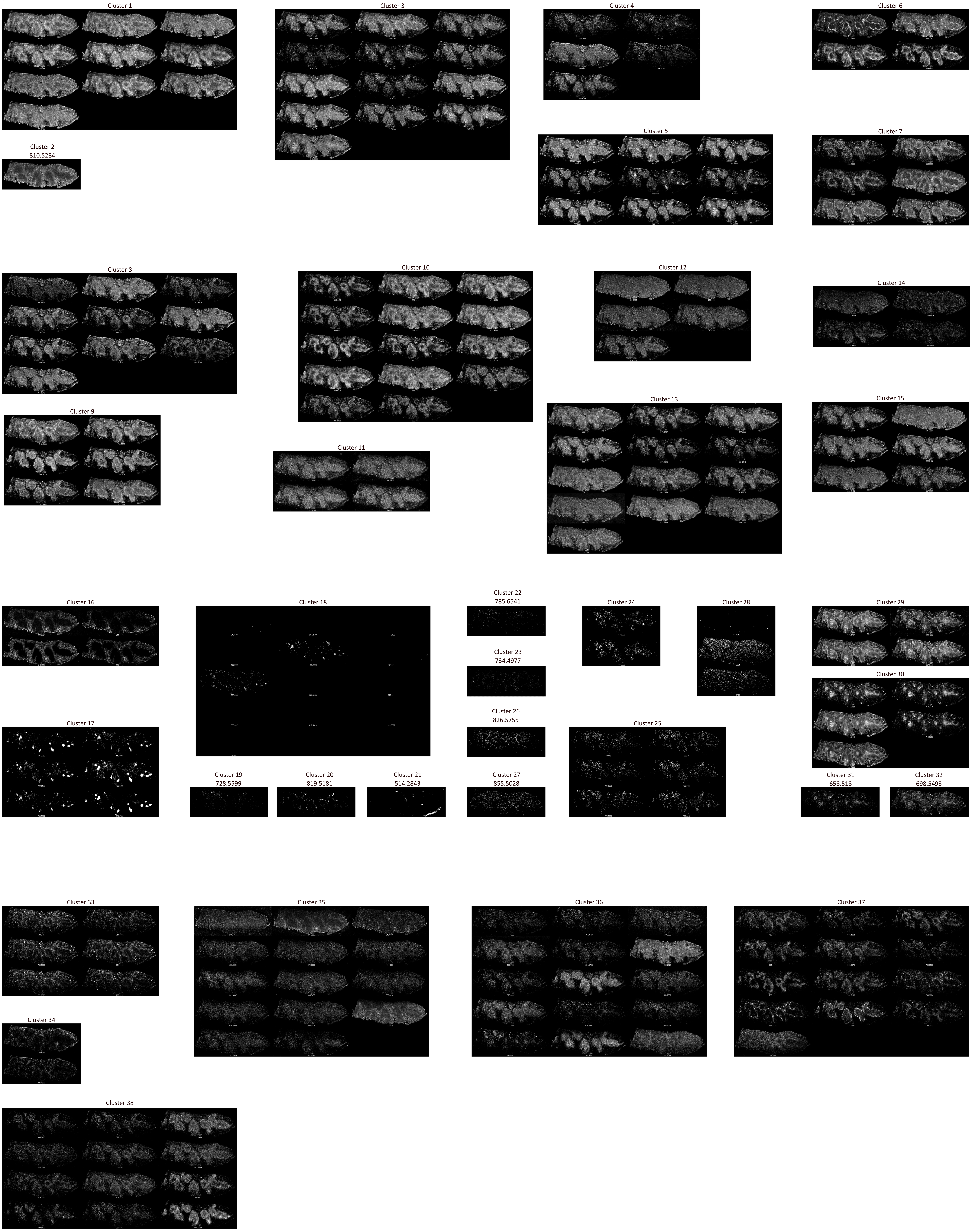
Cluster 1
Cluster 2
810.5284
Cluster 3
Cluster 4
Cluster 5
Cluster 6
Cluster 7
Cluster 8
Cluster 9
Cluster 10
Cluster 11
Cluster 12
Cluster 13
Cluster 14
Cluster 15
Cluster 16
Cluster 17
Cluster 18
Cluster 19
728.5599
Cluster 20
819.5181
Cluster 21
514.2843
Cluster 22
785.6541
Cluster 23
734.4977
Cluster 24
Cluster 25
Cluster 26
826.5755
Cluster 27
855.5028
Cluster 28
Cluster 29
Cluster 30
Cluster 31
658.518
Cluster 32
698.5493
Cluster 33
Cluster 34
Cluster 35
Cluster 36
Cluster 37
Cluster 38
