## Supplemental Figures for "In tissue spatial single-cell metabolomics by coupling mass spectrometry imaging and immunofluorescences"

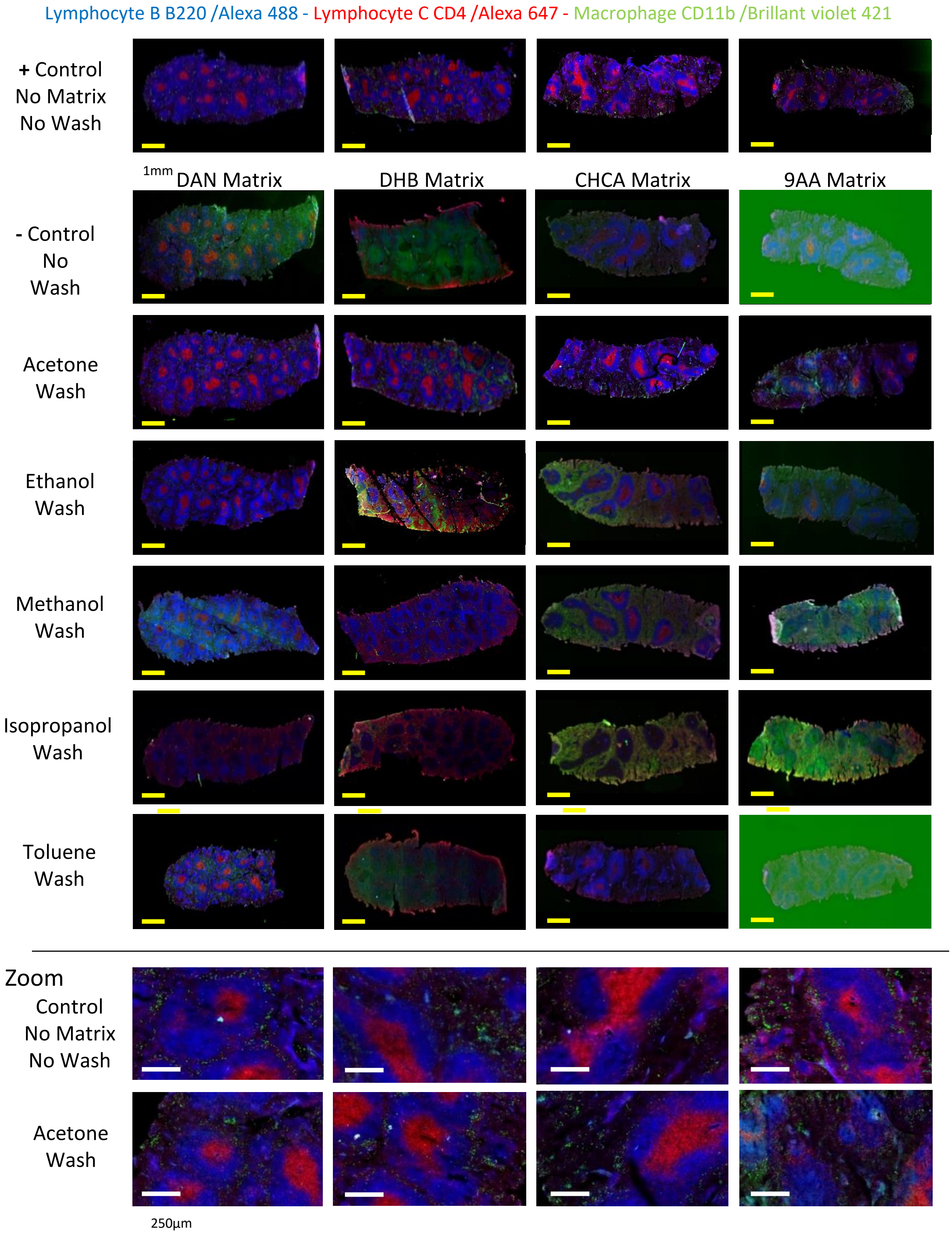


**Figure S1**: Matrix rinsing

Various matrix condition rinsing have been tested. No MSI scans have been performed for this previous optimization. Matrix have been applied according to the Mat & Met section. Slides with matrices deposition have been left one night at room temperature, atmospheric pressure, and in the dark to mime the MSI scan condition in the source. Then, the slides were washed with 2 baths of 2 minutes of the given solvent. Finally, slides are stained for cell receptors B220, CD11b and CD4, following the immunofluorescence protocol described in the Mat & Met section.

Positive control has been performed (untouched condition): after tissue section and thaw-mounting, slides are kept in the freezer waiting to be stained. This condition is made to mime a regular IF staining.

A negative control is a condition with matrix deposition but without any preliminary wash before the IF staining protocol.

### Figure S2 (1/2) : Optimization of the MSI scan condition to limit tissue damage

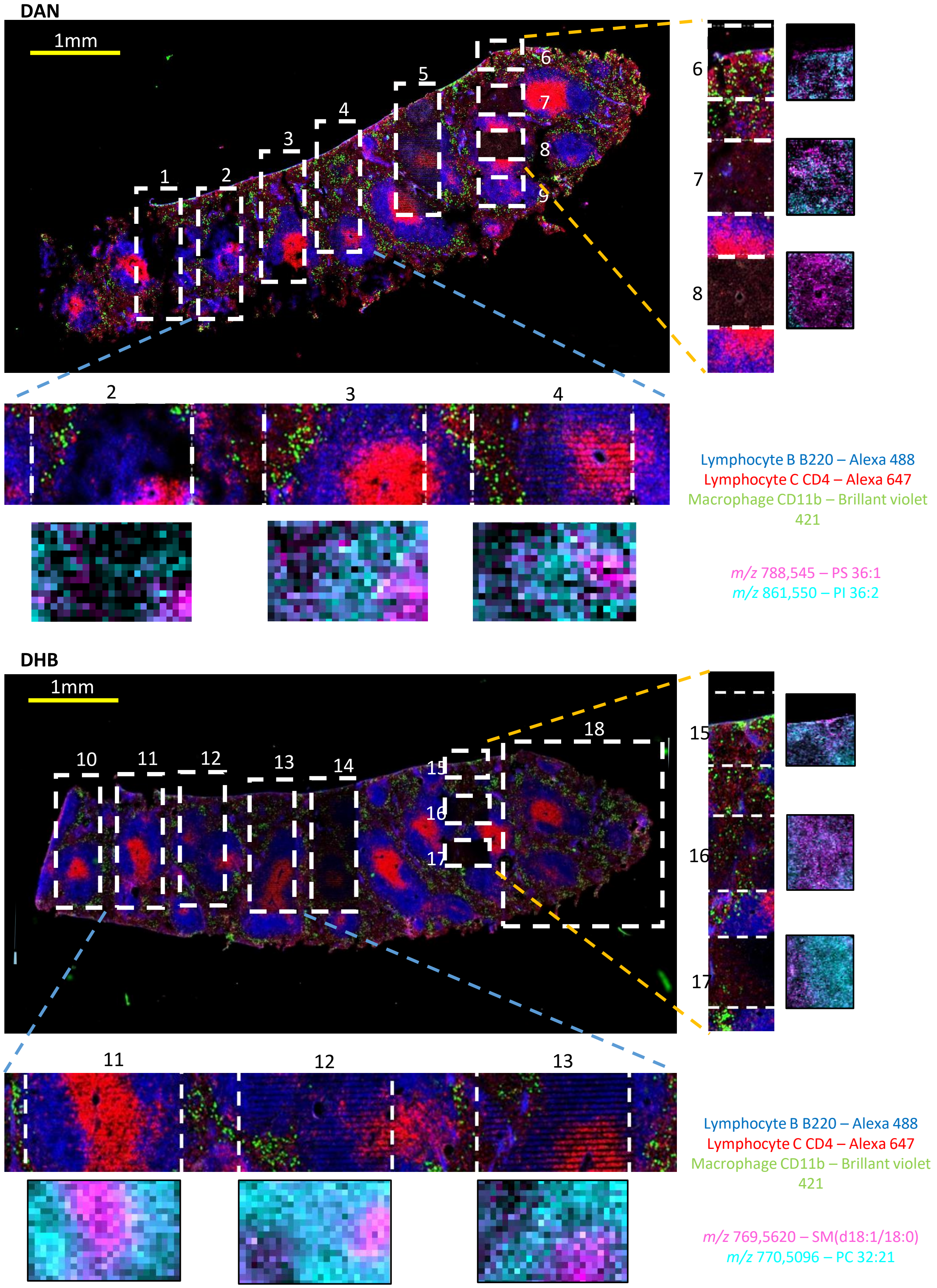

### Figure S2 (2/2) : Optimization of the MSI scan condition to limit tissue damage

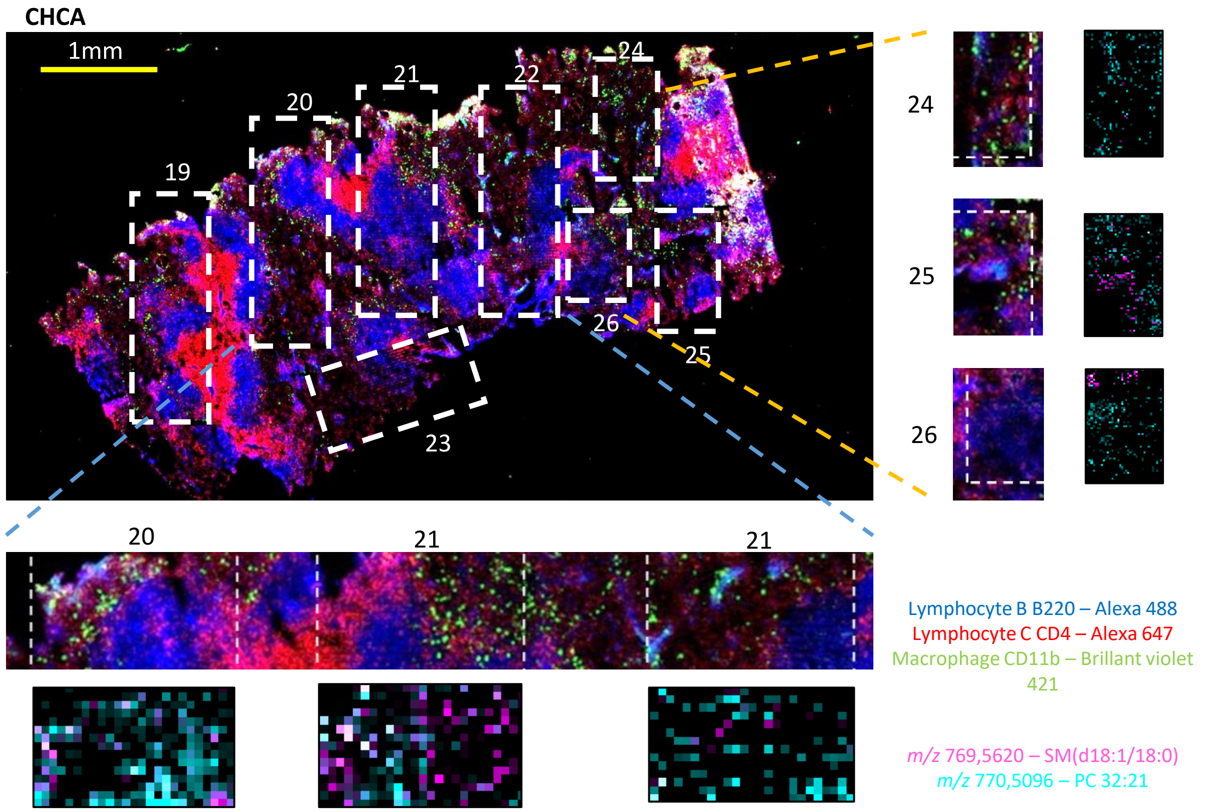


| # ROI | Ionisation Mode | Matrix | Laser energy | | Lateral Resolution | Metaspace (FDR 10%) |
|---|---|---|---|---|---|---|
| | | | Angle | nJ | | |
| 1 | - | DAN | 41 | 6 | 20 µm | **111** |
| 2 | - | DAN | 39 | 7 | 20 µm | **102** |
| 3 | - | DAN | 37 | 18 | 20 µm | **63** |
| 4 | - | DAN | 35 | 38 | 20 µm | **27** |
| 5 | - | DAN | 33 | 109 | 20 µm | **6** |
| 6 | - | DAN | 41 | 6 | 5 µm | **131** |
| 7 | - | DAN | 39 | 7 | 5 µm | **99** |
| 8 | - | DAN | 37 | 18 | 5 µm | **68** |
| 9 | - | DAN | 35 | 38 | 5 µm | **53** |
| 10 | + | DHB | 41 | 6 | 20 µm | **186** |
| 11 | + | DHB | 39 | 7 | 20 µm | **137** |
| 12 | + | DHB | 37 | 18 | 20 µm | **139** |
| 13 | + | DHB | 35 | 38 | 20 µm | **63** |
| 14 | + | DHB | 33 | 109 | 20 µm | **37** |
| 15 | + | DHB | 39 | 6 | 5 µm | **131** |
| 16 | + | DHB | 37 | 7 | 5 µm | **132** |
| 17 | + | DHB | 35 | 18 | 5 µm | **105** |
| 18 | + | DHB | 39 | 7 | 5 µm | **145** |

| # ROI | Ionisation Mode | Matrix | Laser énergy | | Lateral Resolution | Metaspace (FDR 10%) |
|---|---|---|---|---|---|---|
| | | | Angle | nJ | | |
| 19 | + | CHCA | 41 | 6 | 20 µm | **35** |
| 20 | + | CHCA | 39 | 7 | 20 µm | **24** |
| 21 | + | CHCA | 37 | 18 | 20 µm | **18** |
| 22 | + | CHCA | 35 | 38 | 20 µm | **20** |
| 23 | + | CHCA | 33 | 109 | 20 µm | **24** |
| 24 | + | CHCA | 41 | 6 | 5 µm | **22** |
| 25 | + | CHCA | 39 | 7 | 5 µm | **29** |
| 26 | + | CHCA | 37 | 18 | 5 µm | **23** |

**Figure S2:** Optimization of the MSI scanning conditions to limit tissue damage.
Tissue sections are thaw-mounted on a glass slide and the 3 matrices are applied according to the Mat & Met section. Different MSI scan conditions have been performed in negative mode for the DAN matrix and positive mode for DHB and CHCA, with continuous mode (3,7 pixels per second). For each scanned rectangle a different laser energy and lateral resolution condition have been applied according to the joint table. It's important to notice that the AP-SMALDI source Software from TransMIT adjusts the laser fire rate in function of lateral resolution and scan rate to maintain the same amount of shooting per µm.
Once all scans are applied on a tissue, the matrix have been washed with acetone baths (Mat & Met section) prior to IF staining. Tissue damage induced by the different MSI scan conditions can be estimated by comparing it with the rest of the tissue that has not been scanned.
Zoom of the most interested areas are displayed for a better evaluation as well as the corresponding MSI image. To find the best compromised, each MSI-acquired dataset has been uploaded on METASPACE 2020, see here: https://metaspace2020.eu/datasets?prj=aef7411a-97fe-11ed-b433-4f3b39d79cd5&ds_owner=my-datasets
It allowed us to perform an automated annotation and to estimate the quality of each MSI dataset.

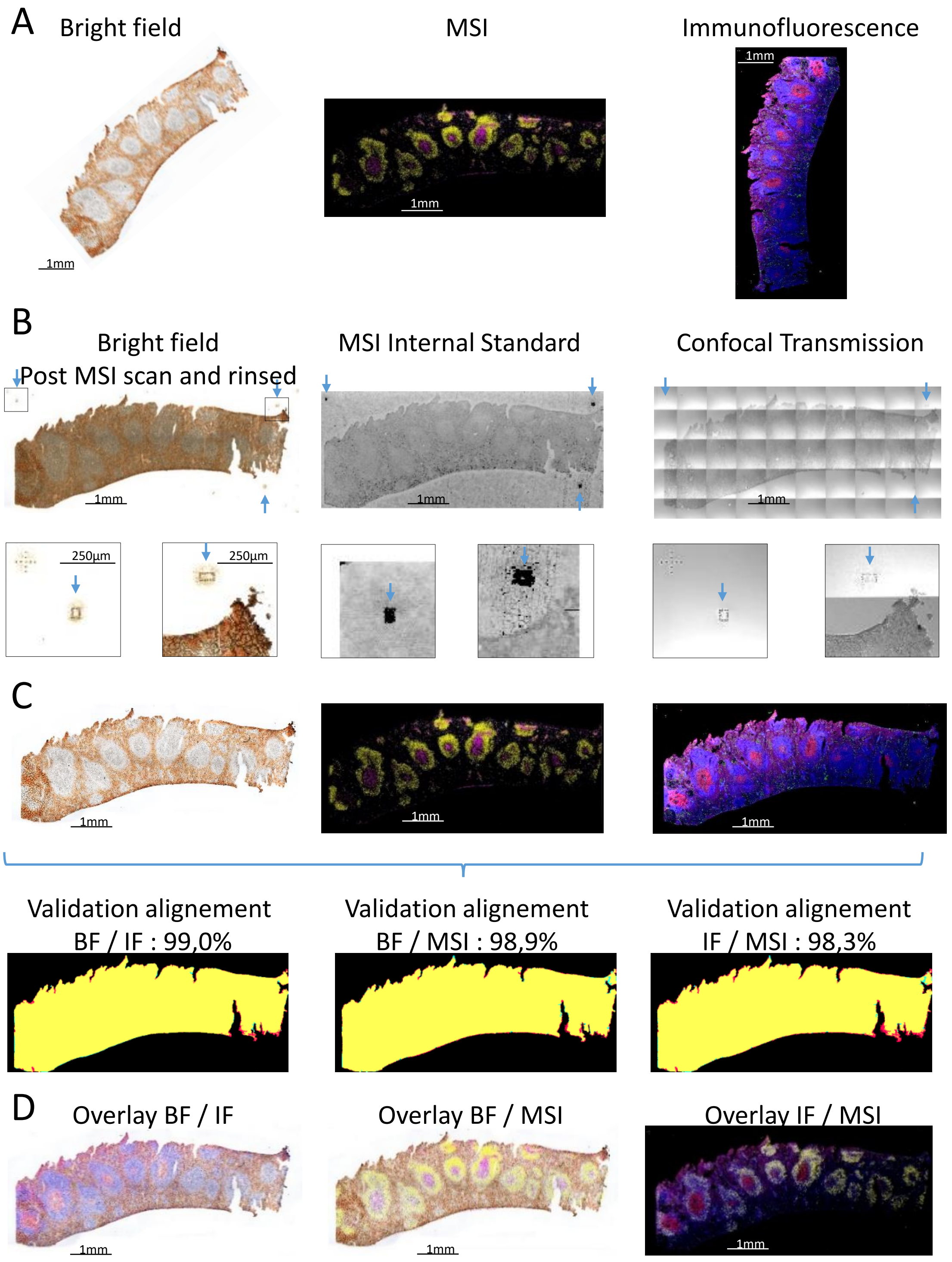


**Figure S3**: Multimodal dataset alignment

a) Display of the bright field image of a tissue section before MSI scanned, associated MSI scan followed by IF staining,

b) Just before MSI acquisition fiducial marks are etched around the tissue with MSI source Laser at full power (see blue arrows). These fiducial marks can be seen on the glass slide in the post-scan bright field image as well as in the transmission image inherently aligned with the IF channel. In MSI datasets, fiducial marks can be seen as imprints, indeed during the etching process MALDI matrix is removed (burnt, destroyed…) and no MSI signal can be obtained from the mark location. By calling matrix peak or internal standard image, locations of fiducial marks can be deduced.

c) Fiducial mark is used to align de different datasets with affine transformation. As the pre-MSI scan bright field image does not present fiducial marks, this image is first aligned with the post-MSI scan one and then they are registered with the other datasets. Once alignment is done quality of registration was assessed using the precision (p), recall (r), and F-score = $(2\times p \times r)/(p+r)$.

d) Results of alignment

imzml file for this figure can be found here: https://metaspace2020.eu/dataset/2022-04-11_11h07m48s

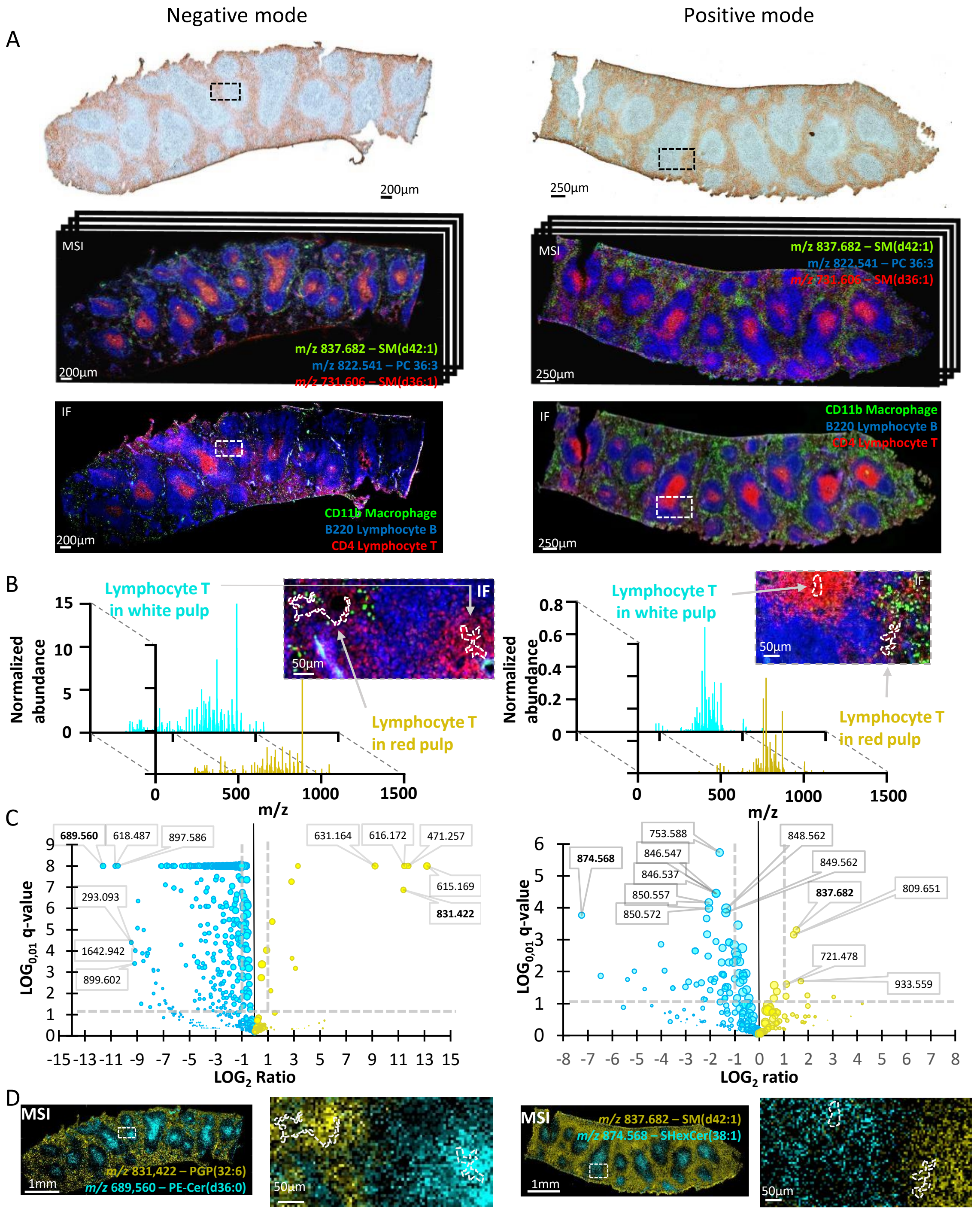


**Figure s4: IF to guide MSI data processing (targeted strategy) – t.test and volcano plot**

On the left side, the spleen section was acquired by MALDI MSI in negative ion mode with the DAN matrix. On the right side, the spleen section was acquired by MALDI MSI in positive ion mode with DHB matrix.

a) Bright field image of spleen tissue sections was acquired before matrix application and MSI scan (top panels). MSI data were acquired in negative and positive mode in linear mode and 5um lateral resolution (middle panels). IF staining for CD11b (green), B220 (blue) and CD4 (red) cells receptor (bottom panels). All multimodal datasets are aligned together with fiducial marks (see **Figure S3**)

b) IF is used to define ROI (in white dashed line), based on an area of lymphocyte T located in white pulp and in Red pulp. Average MS spectra (normalized with internal standard) of the given ROI are then extracted from the MSI dataset.

c) Volcano plot of the statistical univariate analysis performs for each ion. In the x-axis the ratio of ion intensity in Red pulp lymphocyte T area vs white pulp lymphocyte T pulp. In the y-axis, the significance test, here a multiple t-test (FDR approach set à 1% and with Benjamini, Krieger and Yekutieli method) with q value in Log base 0,01 ($Log_{0,01}$(q-value)>1 is considered as significant). The size of the dot correlates with the whole tissue average intensity of corresponding ions. Dash lines indicate the limit of significativity, vertical one for a 2-time ratio and horizontal for a q-value below 0.01.

d) MSI image of significant ions for lymphocyte T in red pulp vs white pulp area with the highest absolute ratio in negative and positive mode.

Ion annotations made with METASPACE 2020, imzml file for this figure can be found here: https://metaspace2020.eu/dataset/2021-11-09_15h54m35s (for négative mode dataset), https://metaspace2020.eu/dataset/2021-11-08_11h28m30s (for positive mode dataset)

| B cells | | | |
|---|---|---|---|
| *m/z* [M-H]$^-$ | Metaspace Annotation | ROC AUC | Ratio (log2) |
| 746.5126 | PE(P-38:6) | 0.8263 | 1.2798 |
| 859.5325 | PI(36:3) | 0.8163 | 0.8831 |

| T cells | | | |
|---|---|---|---|
| *m/z* [M-H]$^-$ | Metaspace Annotation | ROC AUC | Ratio (log2) |
| 701.5119 | PA(36:1) | 0.8653 | 0.9193 |
| 644.5024 | CerP(d36:1) | 0.8649 | 1.9276 |
| 670.518 | CerP(d38:2) | 0.8599 | 2.3512 |
| 794.5699 | PE(40:4) | 0.8591 | 1.2899 |
| 751.5252 | PA(40:4) | 0.8358 | 1.1993 |
| 616.4707 | CerP(d34:1) | 0.8228 | 0.8671 |
| 687.5441 | PE-Cer(d36:1) | 0.8192 | 0.9279 |
| 715.5759 | PE-Cer(d38:1) | 0.8089 | 2.3726 |
| 788.5444 | PS(36:1) | 0.8046 | 0.7867 |
| 630.4863 | CerP(d35:1) | 0.8015 | 1.3402 |

| Red pulp | | | |
|---|---|---|---|
| *m/z* [M-H]$^-$ | Metaspace Annotation | ROC AUC | Ratio ($log_2$) |
| 615.1691 | HEME-$^{56}$Fe | 0.8819 | 4.9119 |
| 306.0764 | Glutathione | 0.8338 | 2.0885 |
| 810.5284 | PS (38:4) | 0.8047 | 0.6986 |

| White pulp | | | |
|---|---|---|---|
| *m/z* [M-H]$^-$ | Metaspace Annotation | ROC AUC | Ratio (log2) |
| 746.5126 | PE(P-38:6) | 0.9106 | 1.3024 |
| 790.5387 | PE(40:6) | 0.8911 | 1.2292 |
| 869.5549 | PI(P-38:4) | 0.8844 | 1.9824 |
| 768.5524 | PE(38:3) | 0.8783 | 1.2614 |
| 751.5252 | PA(40:4) | 0.8770 | 1.2629 |
| 766.5386 | PE(38:4) | 0.8747 | 0.6510 |
| 725.5117 | PA(38:3) | 0.8734 | 0.7666 |
| 794.5699 | PE(40:4) | 0.8676 | 1.3020 |
| 742.5388 | PE(36:2) | 0.8536 | 0.7037 |
| 857.5179 | PI(36:4) | 0.8449 | 0.5135 |
| 792.5539 | PE(40:5) | 0.8394 | 1.3604 |
| 889.5811 | PI(38:2) | 0.8382 | 1.4664 |
| 859.5325 | PI(36:3) | 0.8361 | 0.7011 |
| 714.5078 | PE(34:2) | 0.8348 | 1.1848 |
| 552.2721 | Vignatic acid A | 0.8340 | 1.4069 |
| 748.5284 | PE(P-38:5) | 0.8290 | 0.9518 |
| 749.5105 | PA(40:5) | 0.8279 | 0.7712 |
| 303.2327 | Arachidonic acid | 0.8268 | 1.0007 |
| 863.5629 | PI(36:1) | 0.8259 | 1.0097 |
| 462.2988 | PE(P-18:1) | 0.8158 | 1.1717 |
| 750.5798 | - | 0.8138 | 1.1750 |
| 747.4966 | PA(40:6) | 0.8123 | 0.6884 |
| 794.5341 | PS(P-38:4) | 0.8122 | 1.9710 |
| 909.5492 | PI(40:6) | 0.8119 | 0.9438 |
| 722.5125 | PE P-36:4 | 0.80256 | 0.5855 |

**Table S1: Ions presenting significant ROC AUC for the targeted region of interest in negative mode**
ROC AUC and Ratio ($log_2$) are calculated after normalization with internal standards PE-34:1-d31 [M-D]$^-$ - *m/z* 746,711

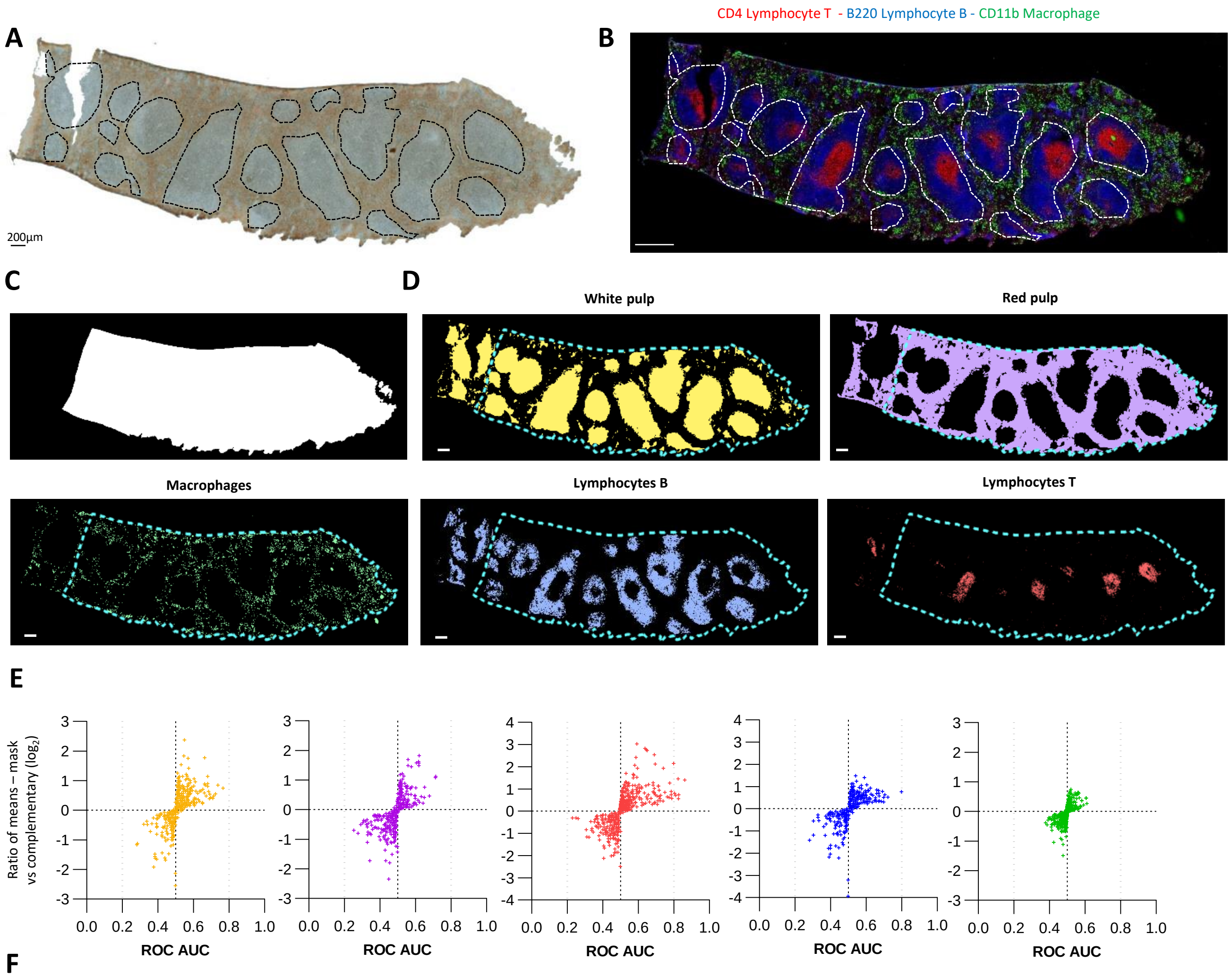


**F**

| *m/z* | Metaspace Annotation | TIC Normalization | | H+ Normalisation | | Na+ Normalisation | | K+ Normalisation | |
|---|---|---|---|---|---|---|---|---|---|
| **B cells** | | ROC AUC | Ratio (log2) | ROC AUC | Ratio (log2) | ROC AUC | Ratio (log2) | ROC AUC | Ratio (log2) |
| 770.5096 | PC 32:1 +$K^+$<br>PE P-38:6 +$Na^+$ | 0.79857 | 0.7646 | 0.74423 | 0.9400 | 0.75582 | 0.9314 | 0.71836 | 0.8111 |

| *m/z* | Metaspace Annotation | TIC Normalization | | $[IS+H]^+$ Normalisation | | $[IS+Na]^+$ Normalisation | | $[IS+K]^+$ Normalisation | |
|---|---|---|---|---|---|---|---|---|---|
| **T cells** | | ROC AUC | Ratio (log2) | ROC AUC | Ratio (log2) | ROC AUC | Ratio (log2) | ROC AUC | Ratio (log2) |
| 769.5618 | SM(d36:1) + $K^+$ | **0.86234** | 1.3976 | 0.80711 | 1.4913 | **0.81424** | 1.4108 | 0.75473 | 1.3560 |
| 848.5565 | PC 38:4 + $K^+$ | **0.82849** | 0.7452 | 0.73258 | 0.8340 | 0.74639 | 0.7474 | 0.71834 | 0.7105 |
| 725.5567 | SM(d34:1) + $Na^+$ | **0.82657** | 0.5681 | 0.75665 | 0.7534 | **0.81828** | 0.7416 | 0.68236 | 0.5170 |
| 753.5885 | SM(d36:1) + $Na^+$ | **0.82379** | 1.4615 | 0.81030 | 1.6539 | **0.83167** | 1.6705 | 0.72950 | 1.3958 |
| 741.5307 | SM(d34:1) + $K^+$ | **0.81154** | 0.7164 | 0.72762 | 0.8378 | 0.73777 | 0.7433 | 0.70606 | 0.6871 |
| 872.5562 | PC 40:6 + $K^+$ (FDR 50%) | 0.79999 | 2.1483 | 0.77723 | 2.1186 | 0.77964 | 2.0550 | 0.73702 | 2.0862 |
| 832.5851 | PC 38:4 + $Na^+$<br>PC 40:7 + $H^+$ | 0.77406 | 0.6196 | 0.75374 | 0.7880 | **0.81931** | 0.7927 | 0.66185 | 0.5645 |
| 731,6063 | SM(d36:1) + $H^+$ | 0.74182 | 1.9296 | 0.75794 | 2.1168 | 0.74673 | 0.7531 | 0.68448 | 1.8538 |
| 810,5988 | PC 38:4 + $H^+$ | 0.72268 | 0.5608 | 0.79833 | 0.6933 | 0.75319 | 0.7531 | 0.63298 | 0.4978 |
| 703.5753 | SM(d34:1) + $H^+$ | 0.71749 | 0.6582 | **0.80706** | 0.8150 | 0.74086 | 0.7869 | 0.63549 | 0.5947 |

**Figure s5: ROC analysis – Metabolic fingerprint in positive mode**

a) Bright-field image of spleen tissue section acquired before MSI analysis in positive mode.
b) IF staining for CD11b (green), B220 (blue) and CD4 (red) cell receptors made after MSI scan.
c) Binary mask of the restricted area where ROC analysis has been performed in order to exclude the damaged area in the left part of the tissue.
d) IF and brightfield images have been used to generate binary masks of the location of the different cell populations and tissue structure: White pulp, Red Pulp, lymphocyte T, lymphocyte B and macrophage. The white scale bar corresponds to 200µm.
e) Volcano ROC (ratio of means = f[ROC AUC]) for the different selected binary masks obtained with TIC normalization.
f) Table of strongest AUC of ROC for each mask with different normalizations: TIC, *m/z* 790.7726-$[M+H]^+$, *m/z* 812.754-$[M+Na]^+$ and *m/z* 827.7397-$[M+K]^+$.
Ion annotations made with METASPACE 2020, imzml file for this figure can be found here: https://metaspace2020.eu/dataset/2021-11-08_11h28m30s

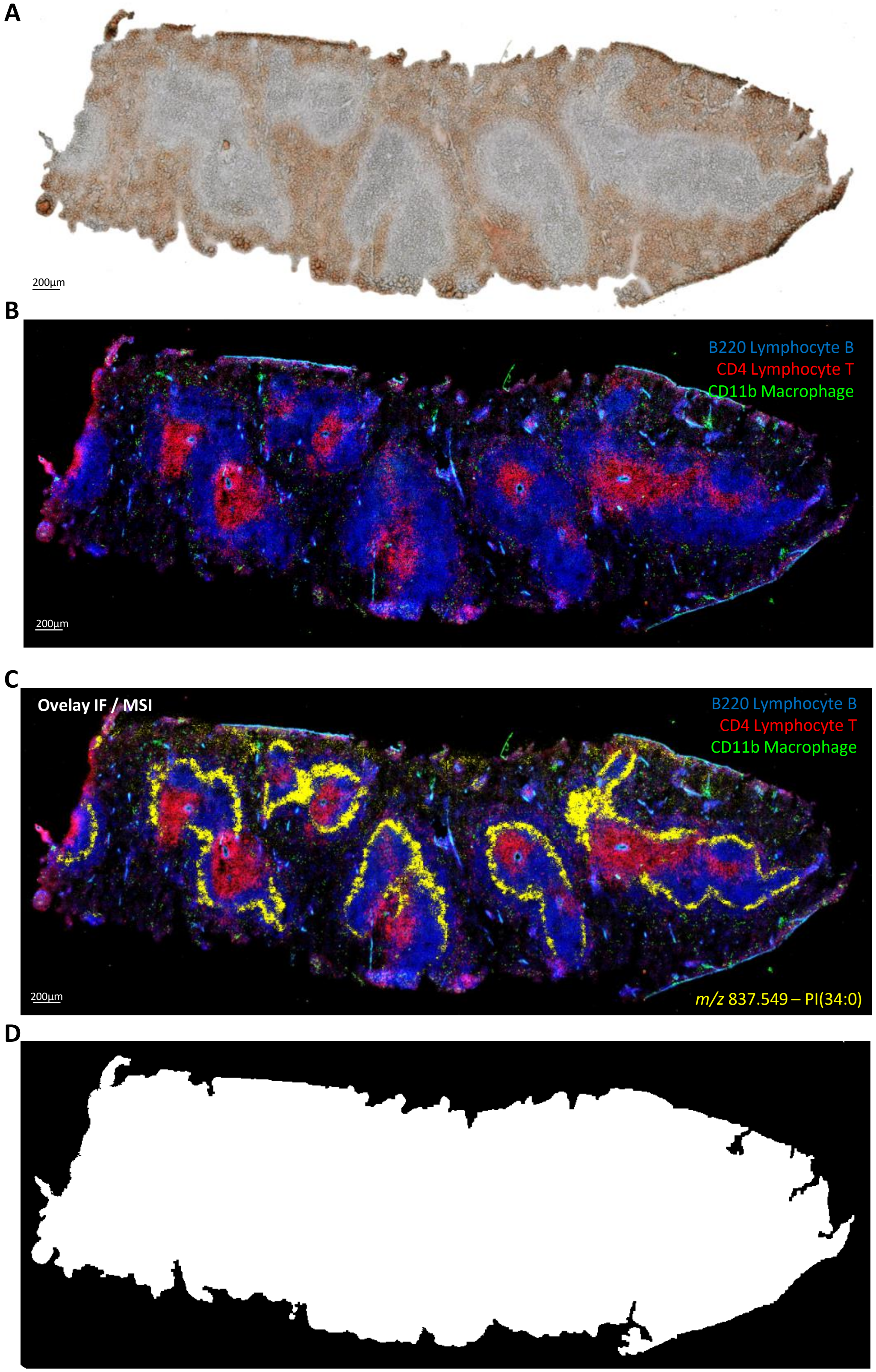


**Figure S6: Histological spleen overview and spatially coherent ion distribution**

a) Bright-field image of mouse spleen section.

b) IF image of spleen section acquired after MSI scan and matrix rinsing.

c) Overlay of IF image and ion map of *m/z* 837.549. https://metaspace2020.eu/annotations?db_id=38&ds=2022-04-15_13h38m32s&mode=Negative&organism=Mus%20musculus%20%28mouse%29&q=RATE&sort=-mz&row=16&norm=true

d) Whole tissue mask for identifying ion distributions IN or OFF sample.

imzml file for this figure can be found here: https://metaspace2020.eu/dataset/2022-04-15_13h38m32s

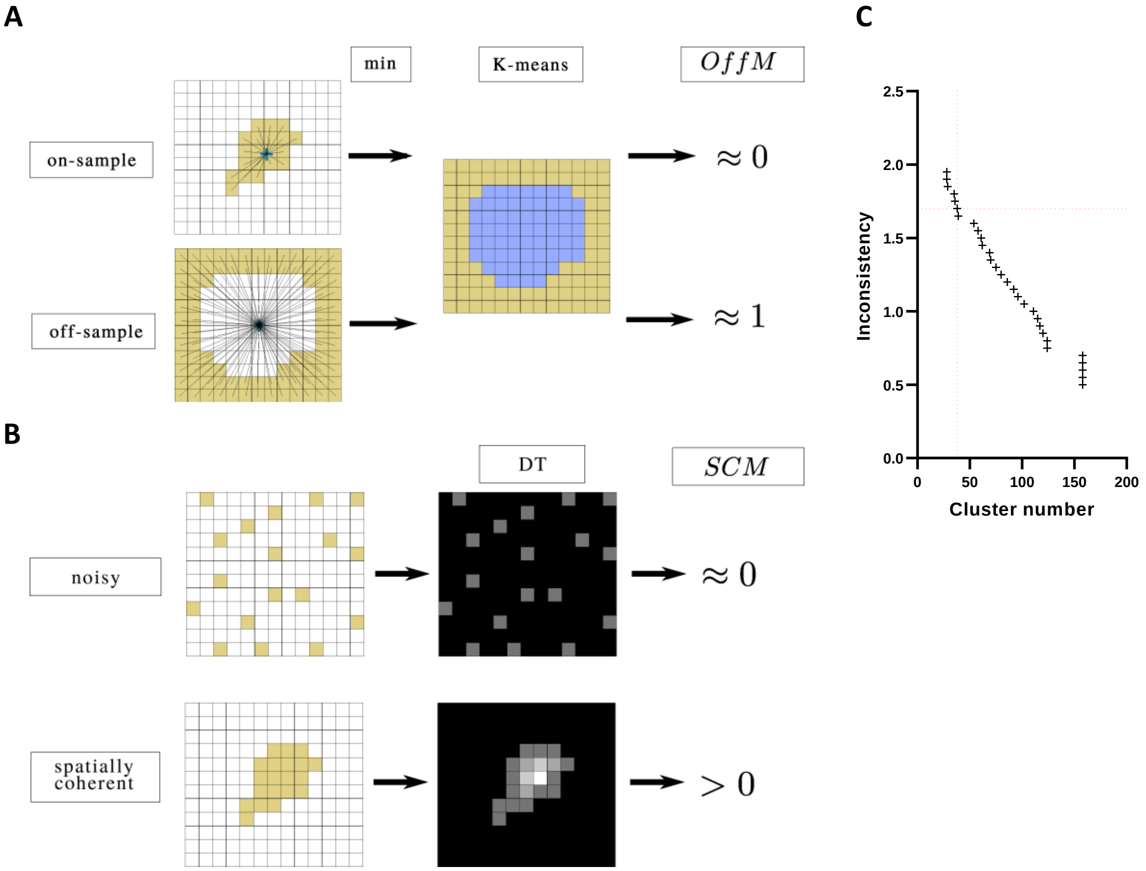


**Figure S7: Schematic of computational methods for the non-guided filtering strategy**

a) Schematic of Off-sample measure computation. The minimum distance between the center and every point in the binarized image is extracted. The distance values are clustered in two groups using the K-means algorithm with k=2. Then, the OffM is computed as the average of the values in the off-sample group (in yellow). The on-sample ions have a value close to 0, while the off-sample ions have a value close to 1.

b) Schematic of Spatially-coherent measure computation. For each ion image, the DT is computed. Then the ratio between the average and the maximum of the DT gives an estimation of the prominence of noise inside the image. SCM values are close to 0 when the ion image is noisy and close to 1 when images are perfectly homogenous and spatially coherent.

c) Inconsistency coefficient plot

**Figure S8: MSI dataset overview - Results of ions filtering and spatial clustering (see attached PDF booklet)**

a) IN and OFF sample peaks filtering: Montage of ion distribution images from different groups. Corresponding m/z can be read at the bottom of each tile (page 2).
b) Spatial coherence sorting: Montages of ions distribution images with high or low spatial coherence score. Corresponding m/z can be read at the bottom of each tile (page 3).
c) Montages of ion distribution images similar or different to IF masks based on ROC AUC. Corresponding m/z can be read at the bottom of each tile (page 3).
d) Average images of the different spatial clusters, associated with the corresponding list of m/z and Metaspace annotation (pages 4-10).
e) KMD projection (base CH2) of relevant m/z grouped by spatial cluster (page 10).
f) Montage of m/z distributions from each spatial cluster. Corresponding m/z can be read at the bottom of each tile (page 11).

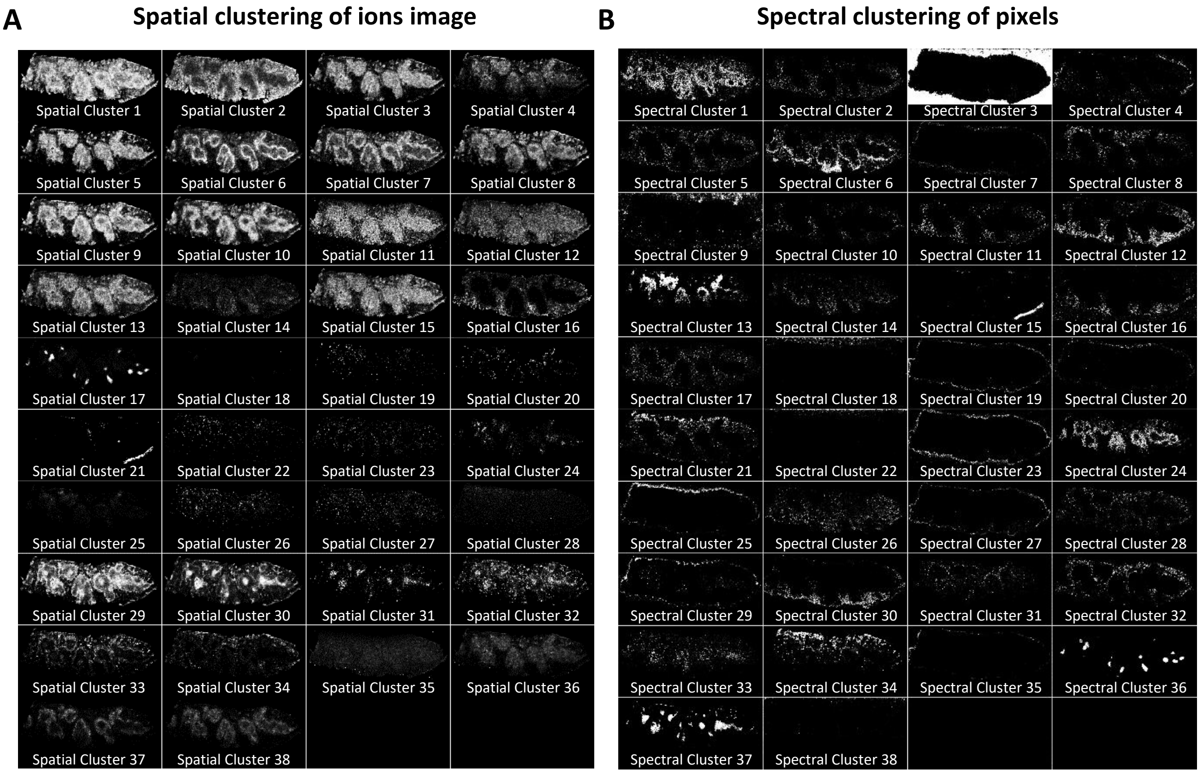


**Figure S9: Spatial clustering vs spectral clustering, complementary strategies to identify different relevant histological areas**

a) Average images of the clusters obtained with spatial hierarchical clustering of *m/z* distributions

b) Clusters obtained with K-means clustering of pixels based on their spectral signature

A

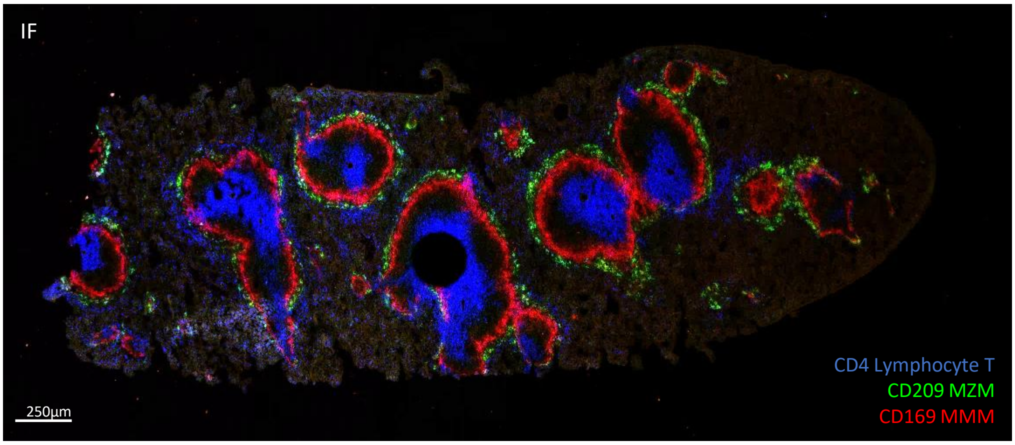


B

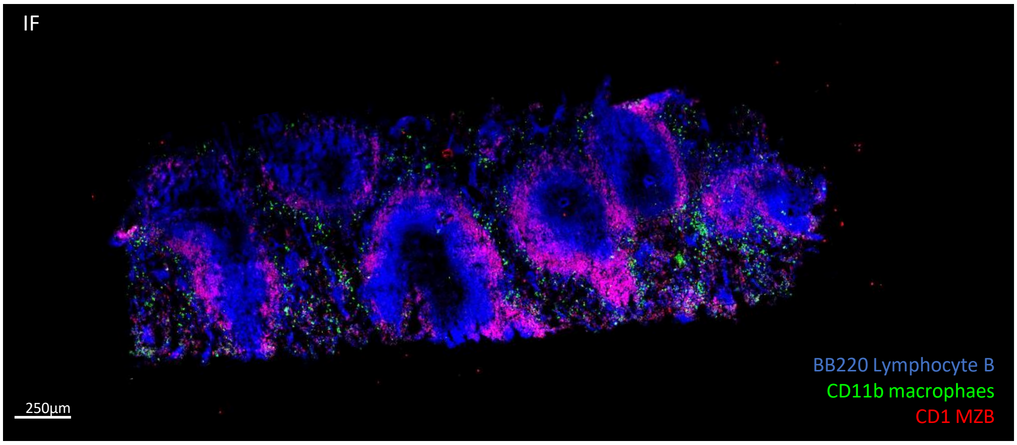


**Figure S10: IF staining of new cell populations based on literature research to match areas highlighted by MSI**
a) IF image of a spleen tissue section stained for CD4, CD209 and CD169 cell receptors
b) IF image of a spleen tissue section stained for B220, CD11b and CD1 cell receptor
These slides have follow a regular IF staining protocol (no matrix spray, no MSI scan prior to immunostaining)

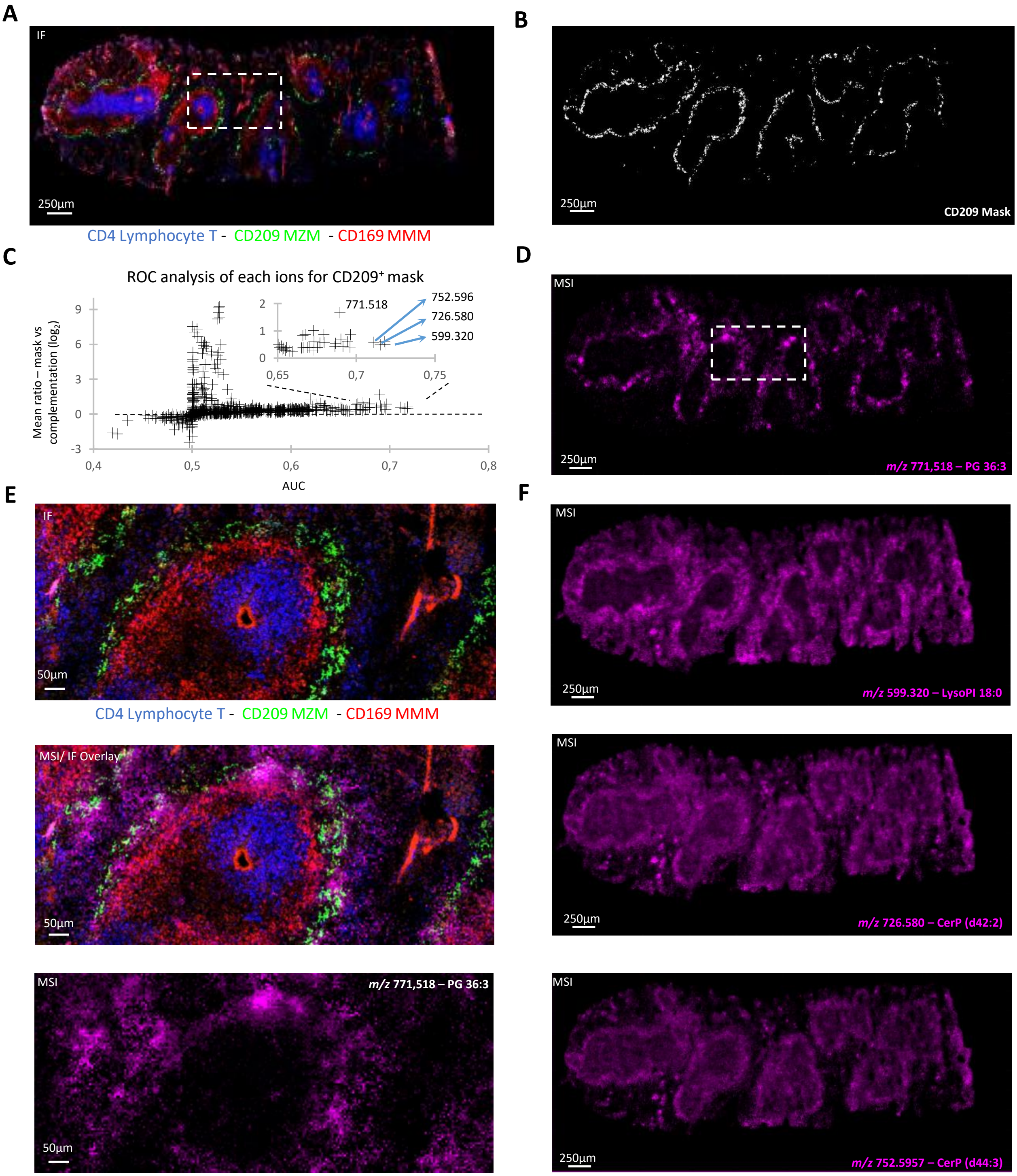


**Figure S11: ROC analysis for CD209 mask**

a) IF spleen tissue section stained for CD4, CD209 and CD169 cell receptors.

b) Display of the mask of CD209+ cells used for ROC analysis.

c) Multiplexed ROC analysis of all detected ions for CD209+ cells mask. The right panel is a zoom on the best classifier ions. None of them present and AUC higher than 0.8, and so cannot be considered as good CD209+ classifier.

d) MSI image of *m/z* 771.518 of spleen tissue section stained for CD4, CD209 and CD169 cell receptors

e) Zoom Overlay of IF image and MSI image *m/z* 771.518.

f) MSI distribution of ROC top 3 ions in the spleen but low ratio.

Ion annotation made with METASPACE, imzml file for this figure can be found here: https://metaspace2020.eu/dataset/2022-04-17_19h49m12s

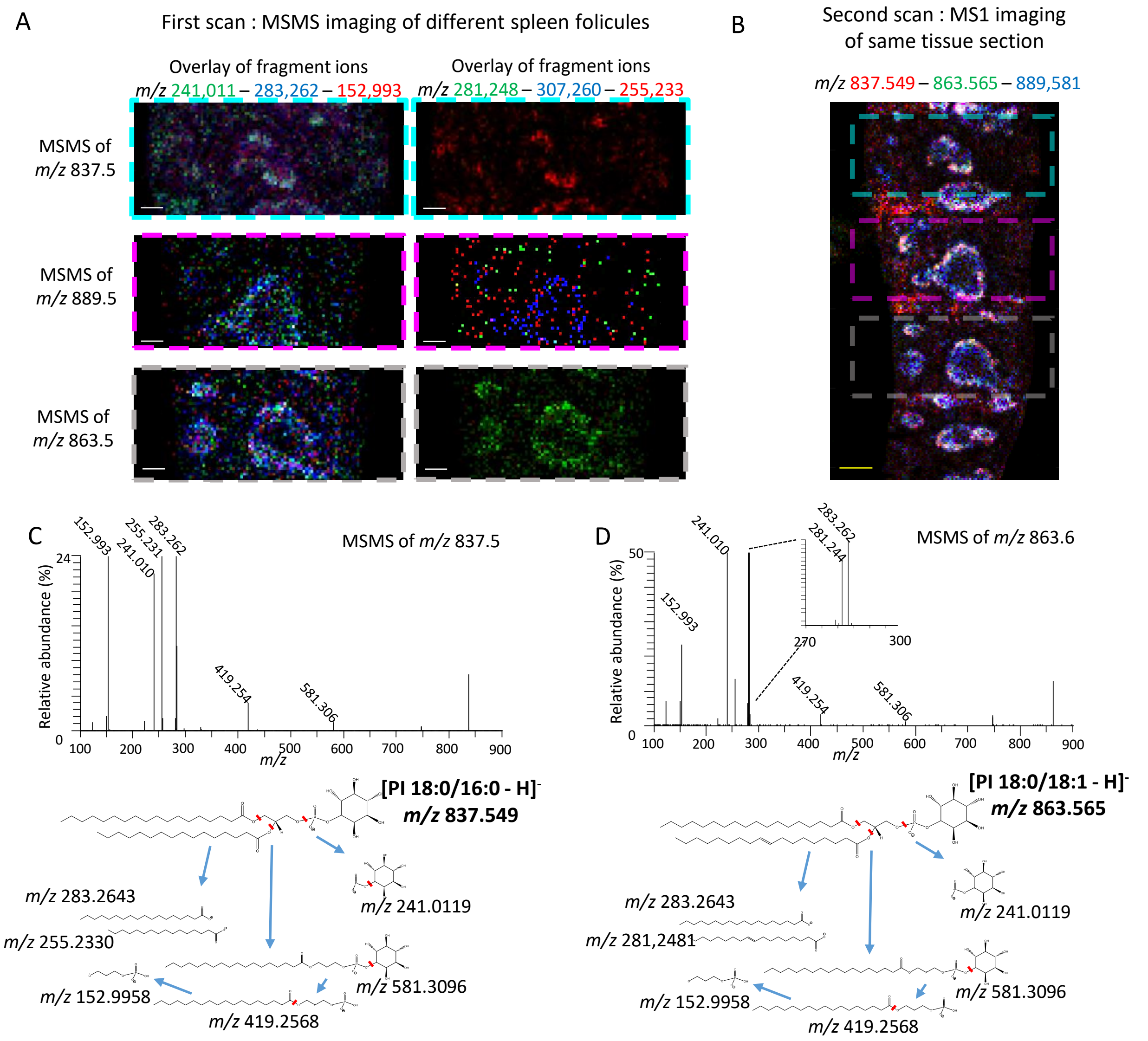


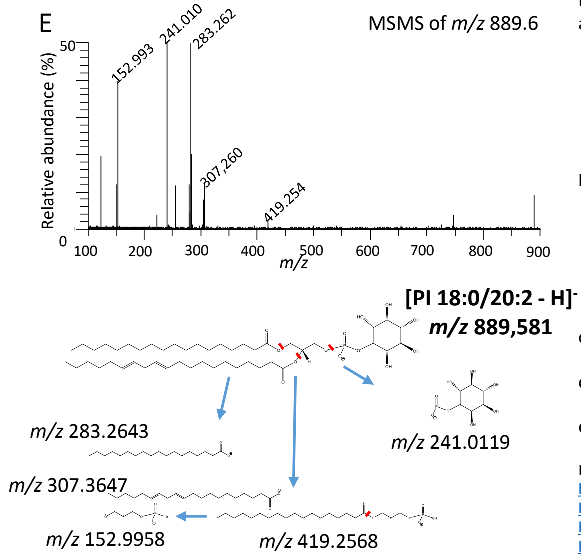


**Figure S12: MSMS Imaging for lipid identifications**

a) MSMS image of parent ion *m/z* 837.5, *m/z* 863.5 and *m/z* 889.5, showing on the **left panel** distribution of *m/z* 152,993 (glycerol phosphate), *m/z* 241,001 (inositol phosphate) and *m/z* 283.262 (stearate). In the **right panel**, images of the specific lipid of each parents' ion are display, *m/z* 255,233 (palmitate), *m/z* 281,248 (oleate), and *m/z* 307.260 (Eicosadienate). MSI scans are perform in negative mode at a lateral resolution of 25µm in full pixel mode. Full pixel mode allows to shoot with a W laser pattern in each pixel in order to recover more signal.

b) MS1 image of the whole's spleen section for the 3 parent ions (no fragmentation in this dataset). Scan areas for the different MSMS images are highlighted in a color dashed rectangle. After MSMS scans, the whole tissue section is rescanned in MS1, in full pixel mode. The remaining matrix present in each pixel is sufficient to obtain an acceptable MS signal. To optimize the MS signal, the scan direction is flipped 90°, so the W scanning pattern of the laser is flipped as well in order to not follow the same path previously scanned in MSMS.

c) MSMS spectra obtained from m/z 837.5 parent and associated fragmentation profile.

d) MSMS spectra obtained from m/z 863,5 parent and associated fragmentation profile.

e) MSMS spectra obtained from m/z 889,5 parent and associated fragmentation profile.

Imzml files for this figure can be found here:
https://metaspace2020.eu/dataset/2022-10-28_14h33m36s (MSMSi of m/z 837.5),
https://metaspace2020.eu/dataset/2022-10-28_14h33m36s (MSMSi of m/z 863.6,
https://metaspace2020.eu/dataset/2022-10-28_14h33m25s (MSMSi ofm/z 889.6),
https://metaspace2020.eu/dataset/2022-10-28_14h33m08s (MS1).
The white scale bar in panel A corresponds to 200µm, Yellow scale bar in panel B corresponds to 500µm.
